# Mammalian metaphase kinetochores are elastic and require condensin for robust structure and function

**DOI:** 10.64898/2025.12.23.696255

**Authors:** Vanna M. Tran, Jinghui Tao, Caleb J. Rux, David Gomez Siu, Miquel Rosas-Salvans, Sophie Dumont

**Author notes:** Corresponding authors (V.M.T.), (S.D.).

## Abstract

Tran et al. show that mammalian metaphase kinetochores undergo large-scale, complex, and elastic deformations under forces of different timescales, magnitudes and directions. They demonstrate that condensin is required for kinetochore local structural organization and microtubule attachment under force.

**Abstract:** The mammalian kinetochore connects chromosomes to dynamic spindle microtubules. To remain attached, it must maintain its structural integrity under force, but to what extent and how it does so remain unclear. Under spindle forces, we find using super resolution microscopy that inner (CENP-A) and outer (Hec1) metaphase kinetochores undergo correlated, large-scale (1 µm) deformations along the force axis, suggesting dynamic, relative sliding of parallel protein linkages. Kinetochore shape changes can be asymmetric, with centromere-facing “tails” correlating with erroneous attachment geometries. Applying microneedle pulling forces, we demonstrate that kinetochores are elastic, stretching under force and relaxing in seconds afterwards. Finally, we show that SMC2 depletion results in more variable kinetochore deformations, despite maintained elasticity, and in reduced microtubule attachment stability. Thus, the kinetochore is structurally highly dynamic and requires a stable centromere base to maintain its structure and function under force. We propose a model whereby individual protein linkages are stiff yet global kinetochore structure flexible to accommodate different attachment geometries and forces while maintaining function.

## Introduction

The kinetochore is a large macromolecular complex essential for maintaining the fidelity of chromosome segregation at cell division. This protein complex is built on the chromatin region defined as the centromere and grips onto spindle microtubules to segregate chromatids. The kinetochore plays a biochemical role ensuring biorientation of all chromosomes before anaphase onset. It also plays a physical role in transmitting and resisting microtubule-generated forces. The mammalian kinetochore has been studied for decades, and we now know much about its molecular architecture and biochemistry (Cheeseman & Desai, 2008; Johnston et al., 2010). We even know from in vitro work the biophysics of how several individual kinetochore proteins bind (Akiyoshi et al., 2010; Demidov et al., 2025; Powers et al., 2009; Volkov et al., 2018). Yet, how the mammalian kinetochore as a whole maintains its structural integrity and function under spindle forces remains poorly understood.

Kinetochore proteins are assembled hierarchically. The most inner, centromeric region on the chromatin is specified by the presence of the histone variant CENP-A. Constitutive Centromere Associated Network (CCAN) proteins that build the inner kinetochore are recruited as the next layer, and outer kinetochore proteins are then recruited to build the layer that binds spindle microtubules (Brinkley & Stubblefield, 1966; Rieder, 1981). This layered structure builds the force transmission pathway along CENP-A to Hec1 linkages that allows chromosome movement, being under tension throughout much of its lifetime (Long et al., 2019; Rago & Cheeseman, 2013; Suzuki et al., 2016). The last two decades have revealed how inner vs outer kinetochore proteins shift position relative to each other along the force axis, within a linkage (Dumont et al., 2012; Suzuki et al., 2014; Wan et al., 2009), and how kinetochore architecture adapts over the course of mitosis (Magidson et al., 2015). However, we know much less about how individual kinetochore protein linkages, bearing load from DNA to microtubules, move relative to their neighbors. For example, we expect inter-linkage movement within the kinetochore to be important to accommodate different microtubule plus-end tip positions (Kiewisz et al., 2022; Leeds et al., 2023; O’Toole et al., 2020) and incorrectly attached kinetochores (Cimini et al., 2001; Khodjakov et al., 1997). How the kinetochore’s material properties allow and constrain any such inter-linkage flexibility, and their underlying basis, remain an open question. Answering it could have implication for kinetochore signal processing (Joglekar & Kukreja, 2017; Liu et al., 2009; O’Connell et al., 2008; Uchida et al., 2009), error correction (Sarangapani & Asbury, 2014), and kinetochore movement, e.g. through microtubule plus-end dynamics coordination (Leeds et al., 2023).

The kinetochore’s architecture of many parallel linkages binding microtubules and transmitting force is thought to be key to its ability to resist force (Rago & Cheeseman, 2013). While protein-protein interactions can break and proteins unfold on the scale of tens to a few hundred pN (Bustamante et al., 2004), metazoan kinetochores are estimated to resist loads of nearly 1nN (Nicklas, 1983; Ye et al., 2016) – with many linkages helping share the load. Global structural integrity between these individual load-bearing linkages could be maintained by proteins that laterally connect individual linkages, individual proteins stretching, and/or through tension diffusing over many parallel attachments (Rago & Cheeseman, 2013). Additionally, since these parallel linkages are all anchored at the centromere, a stiff centromere could help maintain global kinetochore structure. Consistent with this idea, perturbing centromeric chromatin can result in altered kinetochore structure (Ono et al., 2004; Samoshkin et al., 2009) and even reveal its underlying substructures (Kixmoeller et al., 2025; Sacristan et al., 2024; Zinkowski et al., 1991). However, centromeres have been thought of as malleable or plastic structures (Lončarek et al., 2007) and kinetochores as viscoelastic ones (Cojoc et al., 2016). It remains unclear whether these differences are due to measurements from different kinetochore-microtubule attachment modes or mitotic stages, or because these structures have inherently different structural material properties. More broadly, we do not know the role of centromere material properties in mammalian kinetochore material properties and function.

Here, we ask how the mammalian kinetochore maintains its structural integrity under mitotic forces and its underlying basis. To do so, we probe how the kinetochore responds to mechanical force of different magnitudes, geometries, and timescales. We ask this question at metaphase, where the kinetochore is constantly under high forces. We use live-cell super resolution imaging and computational image analysis to define both inner and outer kinetochore shapes. We show that both regions exhibit large-scale (1 µm) deformations consistent with lateral sliding of individual protein linkages within the kinetochore. We mechanically challenge the mammalian kinetochore under altered attachment geometry by exerting exogenous forces using glass microneedles. We find that metaphase kinetochores behave as an elastic material that can not only deform, but can do so in a complex, non-uniform manner, for example with parts of the kinetochore “escaping” from the main kinetochore towards the centromere. Finally, we show through RNAi of condensin subunit SMC2 that centromeric material properties are essential for kinetochore structural maintenance and microtubule attachment under force. We propose that while individual kinetochore linkages are stiff enough to limit their deformation under load and allow local mechanotransduction, global metaphase kinetochore structure is compliant enough to adapt to varied attachment forces and geometries and thus maintain attachment function.

## Results

### Inner and outer mammalian kinetochores actively undergo large-scale correlated shape changes at metaphase

To probe how the mammalian kinetochore maintains its structure under spindle forces, we performed live-cell imaging of both inner (eGFP-CENP-A) and outer (Hec1-HaloTag + Janelia Fluor (JF) 549 dye) kinetochore shapes in metaphase rat kangaroo PtK2 cells. These cells only have 14 chromosomes (Lorenza and Ainsworth, 1972), limiting overlapping kinetochore movements and thus helping analysis of individual kinetochore shapes. We live imaged cells in 4D with an Instant Structured Illumination Microscope (iSIM) (Curd et al., 2015) for its super-resolution ability and low light exposure of cells, and used deconvolution. We reasoned that concurrently imaging both inner and outer kinetochores was essential to constrain the origin of any observed shape changes.

We first sought to measure the length of kinetochores along the spindle’s main force axis, defined here as the sister-to-sister kinetochore axis (K-K). We used TrackMate to follow kinetochores, then summed a 3D image to a 2D image only in the three centermost z-slices, which we then collapsed to a 1D linescan along the K-K axis as proxy for the force axis (Fig. 1 A). Most kinetochores’ intensity linescans (shapes) were well-fit by a Gaussian distribution, with full-width-at-half-max as a metric for size (Fig. 1 B, “Small-scale”; and Video 1). However, some kinetochores took clearly non-Gaussian shapes in both inner and outer kinetochore layers (Fig. 1 B, “Large-scale”; and Video 1). To more accurately measure the lengths of these kinetochores, we used a method that relies on the full intensity linescan information, and not the standard full-width at half maximum (FWHM) method. We used a generalized full width (GFW) method, taking the mean of all lengths under the kinetochore’s intensity linescan – effectively the area under the curve (AUC) divided by the intensity peak height – to measure the length of the kinetochore (Figs. 1 A and B). This GFW method better captures the lengths of non-Gaussian intensity profiles over the FWHM (Fig. 1 B). Using this method, metaphase CENP-A and Hec1 distributions were 435 ± 60 nm and 461 ± 67 nm long along the force axis (Fig. 1 C), with the length averaged for each individual kinetochore over its imaging lifetime. The Hec1 length was significantly longer than CENP-A’s, after accounting for different imaging wavelengths. Since inner and outer kinetochores were imaged in different wavelengths, we confirmed that deconvolution corrected for chromatic aberration using Tetraspeck beads (488 nm beads at 334 ± 14 nm; 561 nm beads at 333 ± 13 nm; Fig. 1 C). Finally, comparison with the measured lengths of 200 nm beads with the above method indicated that kinetochore shapes were much longer than those of a diffraction-limited object (Fig. 1 C).

**Figure 1:**
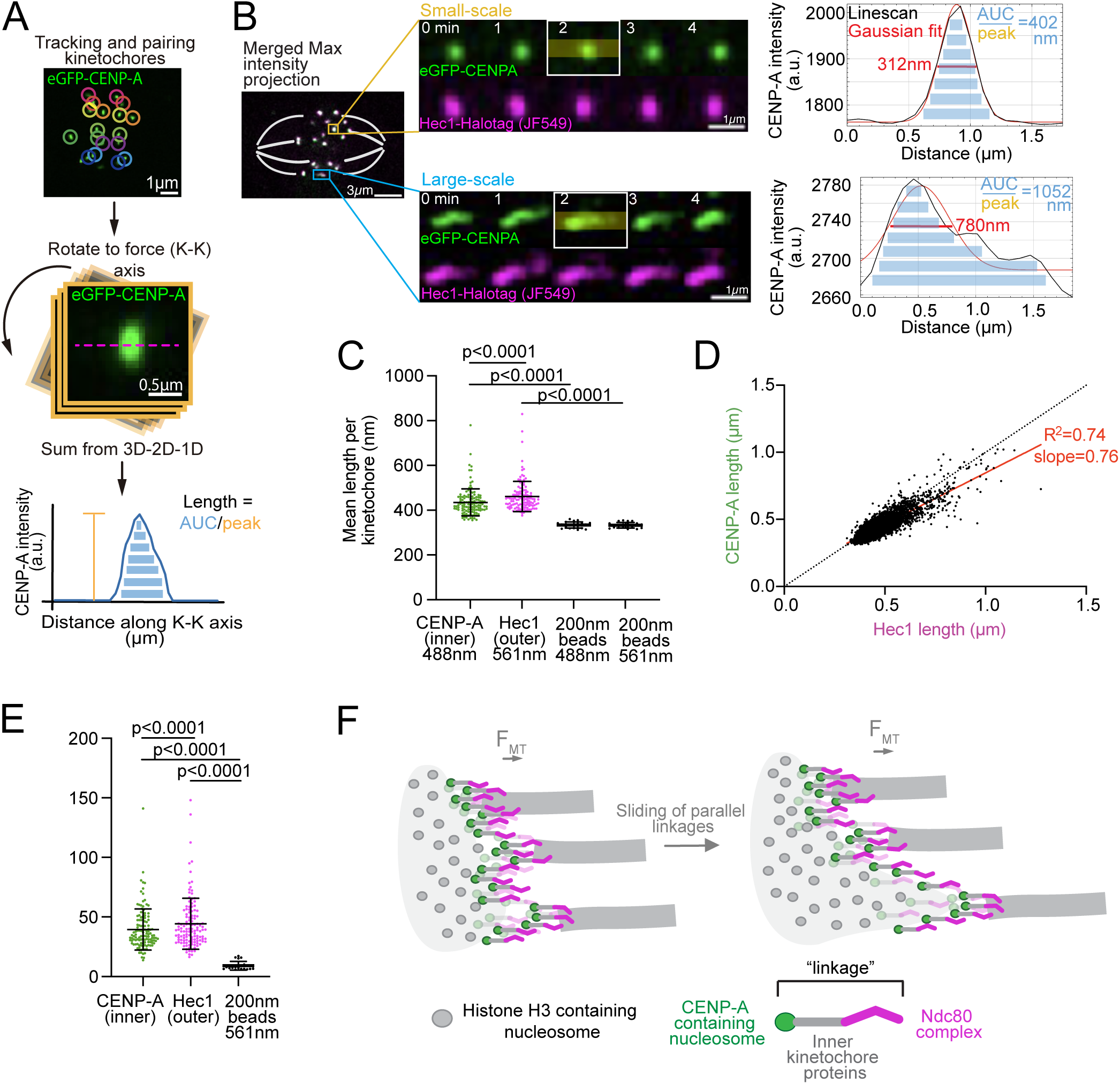
Inner and outer mammalian kinetochores actively undergo large-scale shape changes in a correlated manner at metaphase. (A) Schematic of kinetochore analysis framework. (B) Representative timelapse of PtK2 cell demonstrating a representative small-scale and large-scale deforming kinetochore, visualized by eGFP-CENP-A and Hec1-Halotag with JF549 dye, with intensity linescan along the force (K-K) axis and best Gaussian fits for boxed kinetochores. Red text denotes length measurement using full-width at half maximum of Gaussian fits. Blue text denotes length measurement using AUC/peak calculation. White lines are drawn to approximate spindle shape and k-fibers. (C) Mean length of individual kinetochores over their imaging lifetime for eGFP-CENP-A and Hec1-Halotag with JF549 dye (m = 14 cells, n = 122 kinetochores; mean ± SD; Mann-Whitney test), with control 200 nm beads (in 488 and 561 nm imaging wavelengths) analyzed with the same method (n = 22). **(D)** Correlation between CENP-A length and Hec1 length over each timepoint. Dotted black line indicates a 1:1 comparison and red line indicates the best fit (m = 14 cells, n = 6502 timepoints; Simple linear regression). **(E)** Standard deviation of length of the same individual kinetochores and 200 nm beads as in (C). **(F)** Model cartoon depicting the sliding of parallel linkages that occur to allow for the scale of shape changes observed for CENP-A and Hec1.

In principle, kinetochore shape lengths of > 400 nm could be achieved by individual CENP-A-to-Hec1 linkages sliding with respect to each other and/or stretching of individual CENP-A-to-Hec1 protein linkages (Roscioli et al., 2020). Consistent with inter-linkage movement, there was a strong correlation (R^2^ = 0.74) between CENP-A and Hec1 distribution lengths (simply termed “lengths” thereafter) across all timepoints (Fig. 1 D), indicating a stiff linkage between the inner and outer kinetochore. We also found a strong correlation (R^2^ = 0.90) on a kinetochore-to-kinetochore basis (Fig. S1 A). Also supporting the inter-linkage sliding model, the CenpA-to-Hec1 distance is estimated to be just 107 nm (Wan et al., 2009) and kinetochore lengths reached > 1 µm, suggesting large-scale sliding must occur to account for several hundreds of nanometers of length changes. However, the Hec1 length was longer than CENP-A length by 20-30 nm (Fig. 1 C), consistent with some, though limited, conformational flexibility of the CENP-A to Hec1 linkage.

We then asked if the heterogeneity in kinetochore mean lengths (Fig. 1 C) was due to some kinetochores always being longer or shorter than others, or if some kinetochores actively changed length over time. Consistent with the latter model, inner and outer layers have mean length standard deviations over time of 39 ± 17 nm and 44 ± 22 nm, respectively, while 200 nm beads have significantly lower length standard deviations of 9 ± 4 nm over time (Fig. 1 E). Thus, kinetochores actively change length over time. Finally, we asked if kinetochore width is fixed or if it changes with kinetochore length, as would be expected if kinetochore area or volume were preserved during shape changes. These two different width behaviors have different implications for kinetochore structure and material properties. We binned kinetochores into small- or large-scale length changes (Fig. S1 B) and found that, for kinetochores with large-scale length changes, CENP-A and Hec1 widths (greater than diffraction limited) do not correlate with CENP-A and Hec1 lengths (Figs. S1 C and D). Thus, both inner and outer kinetochore layers actively change length at metaphase in a correlated manner, from diffraction-limited to >1 µm along the force axis without associated kinetochore width changes. Altogether, the data is consistent with a model where relative sliding between parallel linkages gives rise to large-scale kinetochore length changes along the force axis, with sliding magnitude much higher than the CENP-A-to-Hec1 linkage length (Fig. 1 F).

### Erroneous attachment geometries result in complex, asymmetric metaphase kinetochore shapes

Given that some metaphase kinetochores undergo large-scale length changes in both CENP-A and Hec1 (Figs. 1, C, D, and E), we reasoned that some imaged kinetochores may be merotelically attached. We have long known that anaphase kinetochores are elongated when merotelically attached, stuck in the middle of the anaphase spindle while pulled by both spindle halves (Cimini et al., 2001, 2003, 2004; Cojoc et al., 2016; DeLuca et al., 2006; Sen et al., 2021). In metaphase, merotelic kinetochores have also been found to be elongated (Cimini et al., 2004), which we sought to take advantage to define the impact of attachment geometry on kinetochore structure and shape parameters (Fig. 2 A). We enriched for incorrect attachments in PtK2 cells in two ways. First, we exogenously expressed Hec1-9A-HaloTag + JF549 dye which increases the kinetochore’s microtubule affinity and creates hyperstable attachments unable to be resolved by phosphorylation-mediated error correction (Zaytsev et al., 2014). Second, we added AuroraB kinase inhibitor ZM447439 (ZM at 3 µM for 30-90 min) to inhibit error correction (Cimini et al., 2006; Ditchfield et al., 2003). Using iSIM live imaging (Fig. 2 B; and Video 2), we found that in the Hec1-9A condition, outer metaphase kinetochores had increased mean lengths while in the ZM condition, both inner and outer had increased mean lengths (Fig. 2 C). For both, the standard deviations of lengths increased in the inner and outer kinetochore (Fig. 2 D) compared to control. Thus, we confirmed that enriching for erroneous attachments increases the length of metaphase kinetochores.

**Figure 2:**
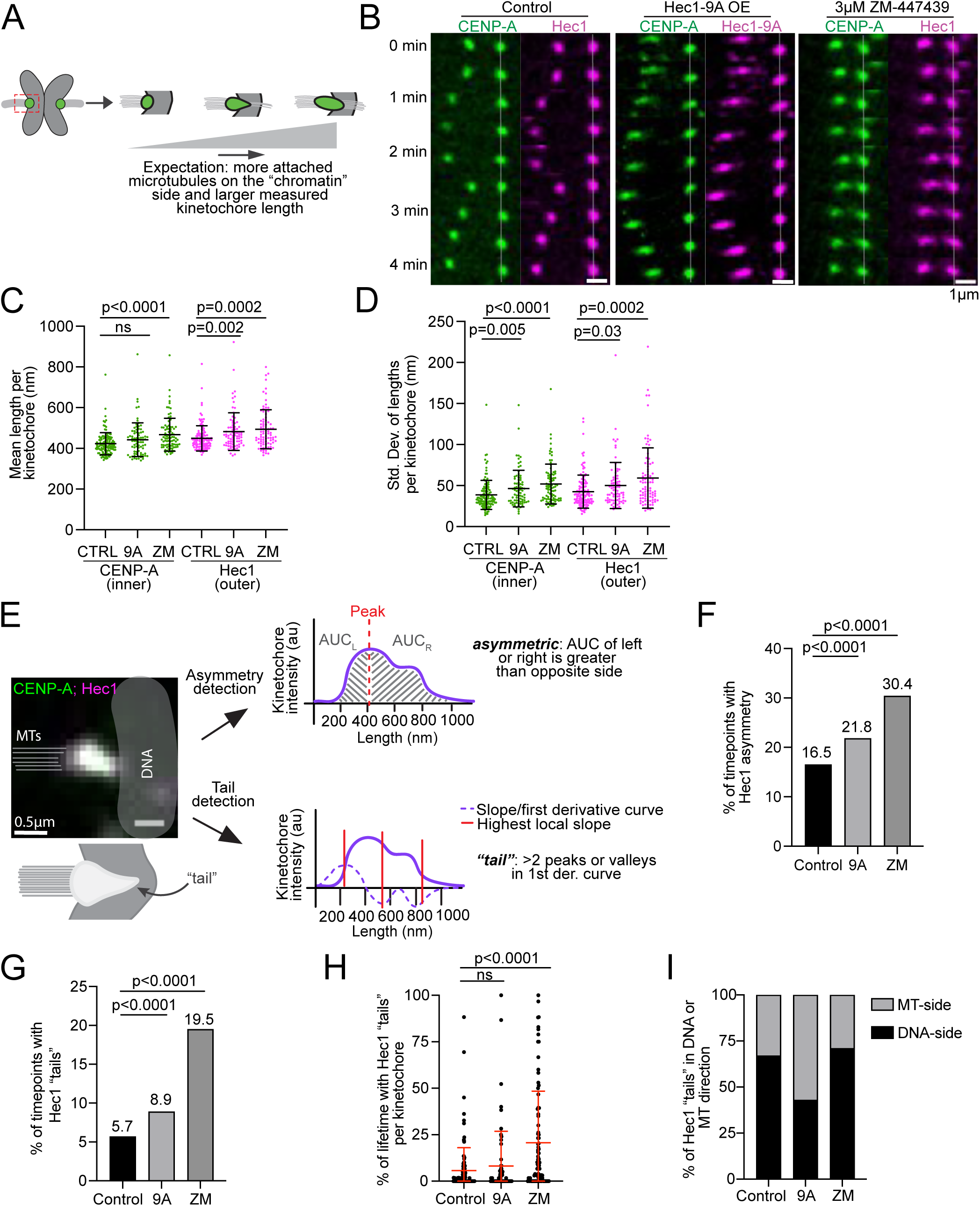
Erroneous attachment geometries result in complex, asymmetric metaphase kinetochore shapes. **(A)** Schematic of how kinetochore shape changes occur with merotelic attachments. **(B)** Representative timelapses of a PtK2 kinetochore pair (eGFP-CENP-A and Hec1-Halotag with JF549 dye) in a control cell, Hec1-9A-HaloTag overexpression cell, and 3 µM ZM447439 (ZM) treated cell, with right kinetochore displayed as stationary (grey line). **(C)** Mean (from left to right columns: 435 ± 60 nm, 442 ± 83 nm, 467 ± 81 nm, 461 ± 67 nm, 482 ± 92 nm, and 494 ± 96 nm) and **(D)** standard deviation (from left to right columns: 39 ± 17 nm, 46 ± 22 nm, 52 ± 24 nm, 44 ± 22 nm, 50 ± 28 nm, 59 ± 37 nm) of individual kinetochores imaging lifetime for eGFP-CENP-A and Hec1-Halotag with JF549 dye control (m = 14 cells, n = 122 kinetochores), Hec1-9A-Halotag (+JF549 dye) transient expression (m = 6 cells, n = 79 kinetochores), and ZM treatment (m = 16 cells, n = 86 kinetochores) (mean ± SD; Mann-Whitney test). **(E)** Cartoon of how kinetochore asymmetry is detected, visualized by CENP-A and Hec1 colocalized (white) with approximated k-fiber microtubules (grey lines, left) and DNA (grey area, right). Cartoon (below) depicts “tail” on the right side of the kinetochore. From the example intensity linescan, asymmetry is detected if the area under the curve (AUC) calculated for left and right side of the peak is greater than opposite side. For tail detection, the first derivative of the intensity curve is calculated. A tail is deemed present if there are >2 minima or maxima. **(F)** Percentage of total timepoints with Hec1 asymmetry detected for control, Hec1-9A expression, and ZM treatment. Significance from Chi-square test. **(G)** Percentage of total timepoints with Hec1 “tails” detected for control (371/6502 timepoints), Hec1-9A expression (280/3131 timepoints), and ZM treatment (929/4762 timepoints). Significance from Chi-square test. **(H)** Percentage of individual kinetochore’s lifetime with Hec1 “tails” detected for kinetochores in (C) and (D) (mean ± SEM; Mann-Whitney test). **(I)** Percentage of tails pointing to microtubule or DNA side for control (n = 222/371 tails towards DNA), Hec1-9A (89/280 tails towards DNA), and ZM-treated (558/929 tails towards DNA) kinetochores, for all individual timepoints with tails.

In both control (Fig. 1 B), Hec1-9A, and ZM (Fig. 2 B; and Video 2) cells, some kinetochores had a non-Gaussian distributed “tail”, with their CENP-A and Hec1 shapes extending asymmetrically either towards the DNA or microtubules. To quantify the frequency of such shapes, we measured the area under the curve (AUC) on each side of the maximum of the kinetochore’s intensity linescan (along the force axis) and compared them (Fig. 2 E). Kinetochore shapes were classified asymmetric if intensities were unbalanced between left and right sides when using the leftmost and rightmost pixel edge of the maximum peak pixel as upper and lower bounds. Here, we used Hec1 to report on shape since we reasoned that if merotelically attached, the outer kinetochore would better report on abnormal kinetochore-microtubule attachment geometries. We observed a significant increase in the fraction of timepoints with Hec1 shape asymmetry in Hec1-9A and ZM compared to control conditions (Fig. 2 F).

To probe what led to these more complex kinetochore shapes, we asked if the “tail” side of the shape distribution was on the DNA or microtubule side. We measured “tail” directions (towards DNA or microtubules) by using the slopes of the first derivative of each kinetochore’s intensity linescan (Fig. 2 E). Kinetochores had “tails” if there were more than 2 local minima or maxima in the first derivative curve (indicating intensity peaks). Similarly to asymmetry, there was a significant increase in Hec1 timepoints with “tails” in Hec1-9A and ZM compared to control (Fig. 2 G). This is consistent with an increase of kinetochores with an unbalanced merotelic configuration (Cimini et al., 2003; Sen et al., 2021), i.e. with different numbers of microtubules bound to each pole. For both Hec1-9A and ZM, kinetochores spend more of their imaging lifetime with “tails” compared to control (Fig. 2 H). For the majority of “tail-forming” kinetochores, the “tails” pointed towards the DNA rather than microtubules, consistent with a few (rather than a full complement of) microtubules being attached to the opposite spindle pole (Figs. 2, A and I). Interestingly, the Hec1-9A condition led to the increased frequency of “tails” facing the microtubule side (Fig. 2 I). Such microtubule-facing tails could, for example, reflect merotelic attachments where there are more microtubules bound on the incorrect than correct spindle side. Together, the data show that increasing incorrect attachments increases both the asymmetry and “tail” formation of metaphase kinetochore shapes, with most “tails” pointing toward the DNA. This suggests that some of the control metaphase kinetochore with “tails” (Fig. 1 B) are merotelically attached and that quantitative kinetochore shape analysis could be used as a reporter of metaphase attachment geometry in live cells.

### Acute force application reveals elastic mammalian kinetochores at metaphase

Kinetochore material properties ultimately underlie their shapes and functions, for example during attachment formation and correction. To probe the material properties of metaphase kinetochores, we asked how force, from both endogenous and exogenous sources, impacts their shape. We measured K-K sister kinetochore distance (K-K distance) as a proxy for force on the kinetochore (Fig. 3 A). As K-K distance increased under endogenous spindle forces during metaphase oscillations, kinetochore lengths also increased (Fig. 3 B, black line). We then asked if all kinetochores stretch with force, or only those that more or less actively change shape, determined by the range of lengths a kinetochore takes over metaphase (Fig. S1 B).

**Figure 3:**
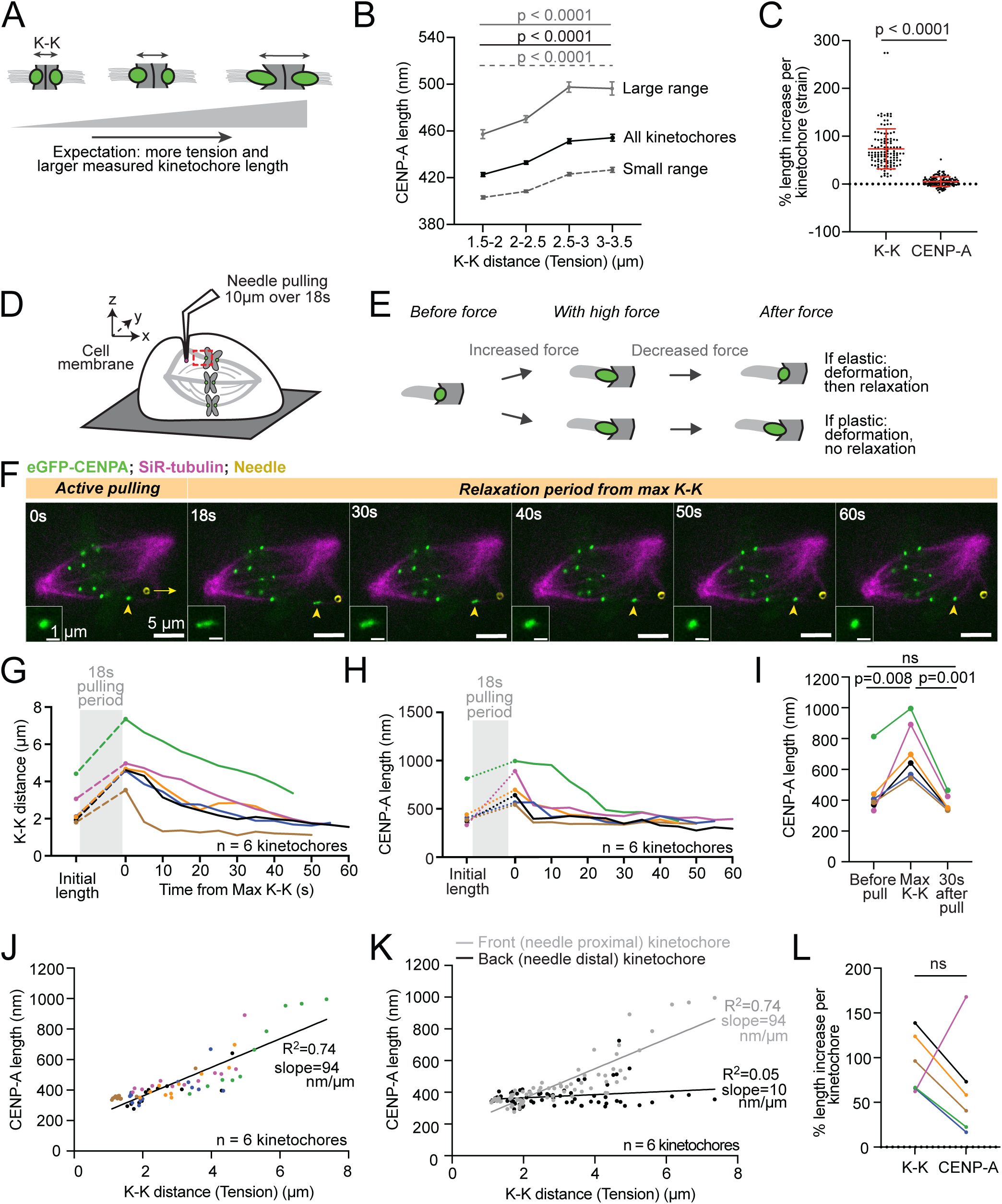
Acute force application reveals elastic mammalian kinetochores at metaphase. (A) Schematic of how kinetochore length changes occur with tension. **(B)** All (black line; m = 122 kinetochores; timepoints per increasing K-K bin n = 1420, 2623, 1518, 617), large-range (grey continuous line; m = 48 kinetochores; per K-K bin n = 522, 1054, 580, 246), and small-range (grey dashed line; m = 74 kinetochores; per K-K bin n = 898, 1569, 938, 371) kinetochores’ Hec1 lengths binned by K-K distance (tension), with bins with < 100 timepoints excluded. Large-range kinetochores were assigned by range of Hec1 lengths over each kinetochore’s imaging lifetime > mean range of all kinetochores. Small-range kinetochores have length ranges < mean range of all kinetochores (mean ± SEM; Mann-Whitney test done for first and last K-K distance bins). **(C)** Percent of K-K distance and CENP-A length increase per kinetochore during imaging lifetime; 74 ± 42% and 5 ± 10% (mean ± SD; Paired t-test). **(D)** Cartoon depicting assay to exert controlled force (10 µm over 18 s) on a k-fiber and its attached kinetochore with a glass microneedle, without membrane rupture (Long et al., 2020; Suresh et al., 2020). **(E)** Potential outcomes of microneedle pulling before force, with high force, and after force: attached kinetochore deforms then relaxes (elastic response) or deforms and does not relax (plastic response). **(F)** Representative PtK2 cell timelapse of eGFP-CENP-A deforming with the microneedle pulling, and relaxing. The yellow arrowhead denotes the “front” sister kinetochore and the long yellow arrow indicates pulling. **(G)** K-K distance over time before, during, and after microneedle pulling for 6 kinetochores. “Initial” length is the frame before pulling begins. Time = 0 s corresponds to the time maximum K-K distance was measured during pulling. The grey box indicates the 18 s pulling period. **(H)** eGFP-CENP-A lengths over time before, during, and after microneedle pulling for 6 kinetochores as defined in (G). The grey box indicates the pulling period, during which kinetochore lengths were not measured due to motion blur. **(I)** CENP-A length before microneedle pulling (∼5 s before), at the maximum K-K distance during pulling, and 30 s after pulling. N = 6 kinetochores (paired t-test). **(J)** Correlation of CENP-A length and K-K distance during the relaxation period after pulling (n = 6 kinetochores). R^2^ = 0.74, significantly non-zero using simple linear regression. **(K)** Correlation of “back” (needle distal) kinetochore CENP-A length (black dots) and K-K distance during the relaxation period after pulling overlaid on “front” (needle proximal) kinetochore lengths (grey dots) as in (I). R^2^ = 0.05 for “back” kinetochores, not significant from zero using simple linear regression. **(L)** Length fold change (strain) of CENP-A and K-K distance (92 ± 33%) and CENP-A (63 ± 56%) from before pulling and at maximum measured K-K distance (n = 6 kinetochores; Paired t-test).

Both groups of kinetochores adopted longer shapes under higher forces (Fig. 3 B): the majority of kinetochores that have small deformations (small range), likely encompassing mostly correct attachments, and those that have large deformations (large range), likely including incorrect attachments (Fig. 2 C). Further, examining kinetochores undergoing regular metaphase oscillations with persistent directional movements, and thus likely correctly attached (Sen et al., 2021), kinetochore lengths of both sisters peaked when one sister switched from poleward to antipoleward movement (P-to-AP) – the exact moment force (and K-K distance) peaks on a metaphase kinetochore (Dumont et al., 2012; Wan et al., 2012) (Fig. S2 A). Since both sister kinetochores change plus-end dynamics states at different times (VandenBeldt et al., 2006; Wan et al., 2012) and in different directions, yet both peak in length at the same time, this kinetochore lengthening likely results from force changes rather than from microtubule dynamics changes. Together, these findings are consistent with kinetochores in both correct and incorrect attachment states deforming under endogenous mechanical force.

To probe the material properties of metaphase kinetochores, we first sought to compare their stiffness to that of centromeres. To do this, we calculated the approximate “strain” for each kinetochore and centromere under the same force using CENP-A length and K-K distance changes, respectively. For each centromere, we used the percent increase of each kinetochore pair’s average three lowest and average three highest K-K distance lengths during its imaging lifetime. For each kinetochore, we used the corresponding timeframes and calculated the percent increase in CENP-A lengths during that same period. We found that kinetochores deformed much less than the centromere under endogenous spindle forces (Fig. 3 C). This suggests that the kinetochore is stiffer than the centromere it is built on, and that its increased stiffness stems from kinetochore-specific elements.

To more directly probe kinetochore material properties, we used microneedle manipulation to directly apply acute, external, and controlled force on kinetochores in Ptk2 cells. We applied exogenous force over the seconds timescale rather than the minutes timescale of endogenous metaphase oscillation forces. A strong, rapid pull is essential toward exerting force on the kinetochore since a weaker or slower force can be dissipated by microtubule polymerization (Long et al., 2020). We first placed the microneedle in the cell next to an outer kinetochore-microtubule bundle (k-fiber, marked by SiR-tubulin), with the membrane deforming around the needle without rupture (Long et al., 2020; Suresh et al., 2020) (Fig. 3 D). To apply force on the attached kinetochore, we pulled the k-fiber perpendicular to the kinetochore-pole axis for 10 µm over 18 s then held the needle in place (Rosas-Salvans et al., 2025). We used spinning disk confocal microscopy to image the shapes of eGFP-CENP-A-labelled kinetochores before, during, and after the pull (Figs. 3, D, E and F; and Video 3). We measured the K-K distance throughout as a proxy for force on the kinetochore. The maximum K-K distance during pulling was higher than without pulling, giving us confidence that pulling exerted a higher force than control (Fig. 3 G compared to Fig. 3 B).

To probe whether metaphase kinetochores behave as elastic or plastic materials (Fig. 3 E), we examined the length of needle-proximal (“front”) kinetochores with a length increase of >100 nm under microneedle pull. Lengthening during pulling followed by relaxation to baseline length post-pulling would indicate an elastic kinetochore. In turn, lengthening with no or incomplete relaxation would indicate a plastic kinetochore. While the mammalian centromere has been reported to be plastic (Lončarek et al., 2007), anaphase merotelic kinetochores have been reported to have both elastic and plastic components (Cojoc et al., 2016). We measured CENP-A length changes before pulling, at the maximum measured K-K, and after microneedle pulling, but not during the pull due to motion blur during needle stepping. We found that deforming kinetochores relax back to their baseline lengths in seconds, consistent with metaphase kinetochores deforming elastically (Figs. 3, E, H, and I; and Video 3) and similar to anaphase merotelics (Cojoc et al., 2016). There was a significant correlation between CENP-A length and K-K distance or force (Fig. 3 J), consistent with deformations observed under endogenous spindle forces (Fig. 3 B). While the needle-proximal “front” kinetochore attached to the pulled k-fiber deformed under acute microneedle force, the needle-distal “back” sister kinetochore typically appears to not deform (Fig. 3 K), suggesting rapid force dissipation between the “front” and “back” kinetochores. While under endogenous, minutes-long forces kinetochore strain was much lower than centromere strain (Fig. 3 C), both strains were similar under these stronger, seconds-long microneedle forces (Fig. 3 L). This could reflect kinetochore strain-softening or centromere strain-stiffening under higher exogenous forces, or timescale-dependent material properties of either the centromere or kinetochore. Together, these findings support a model whereby the mammalian metaphase kinetochore is elastic, able to undergo large-scale (>1 µm) deformations under acute force and to relax back in seconds without permanent change to its shape.

### Loss of condensin leads to variable metaphase kinetochore shape and loss of structural stability

Fragmented and structurally defective kinetochores can result in errors during division (Zielinska et al., 2019). Given that erroneous attachments (Fig. 2) and acute force application (Fig. 3) result in large-scale metaphase kinetochore shape changes, we asked what mechanisms constrain these deformations. Since both Hec1 and CENP-A kinetochore shapes deform in a highly correlated manner (Fig. 1 D), we hypothesized that the base of the kinetochore, inward of CENP-A, constrained these deformations. The kinetochore is assembled on chromatin, and as such we tested the role of SMC2, a subunit of both condensin I and II complexes essential for chromatin compaction (Fig. 4 A). We designed and tested SMC2 siRNA sequences in rat kangaroo cells. We verified effectiveness of SMC2 RNAi after 48 h by qPCR (Fig. 4 B) and by an increased K-K distance (Fig. 4 C), expected from chromatin decompaction (Gerlich et al., 2006; Ribeiro et al., 2009; Sun et al., 2018).

**Figure 4:**
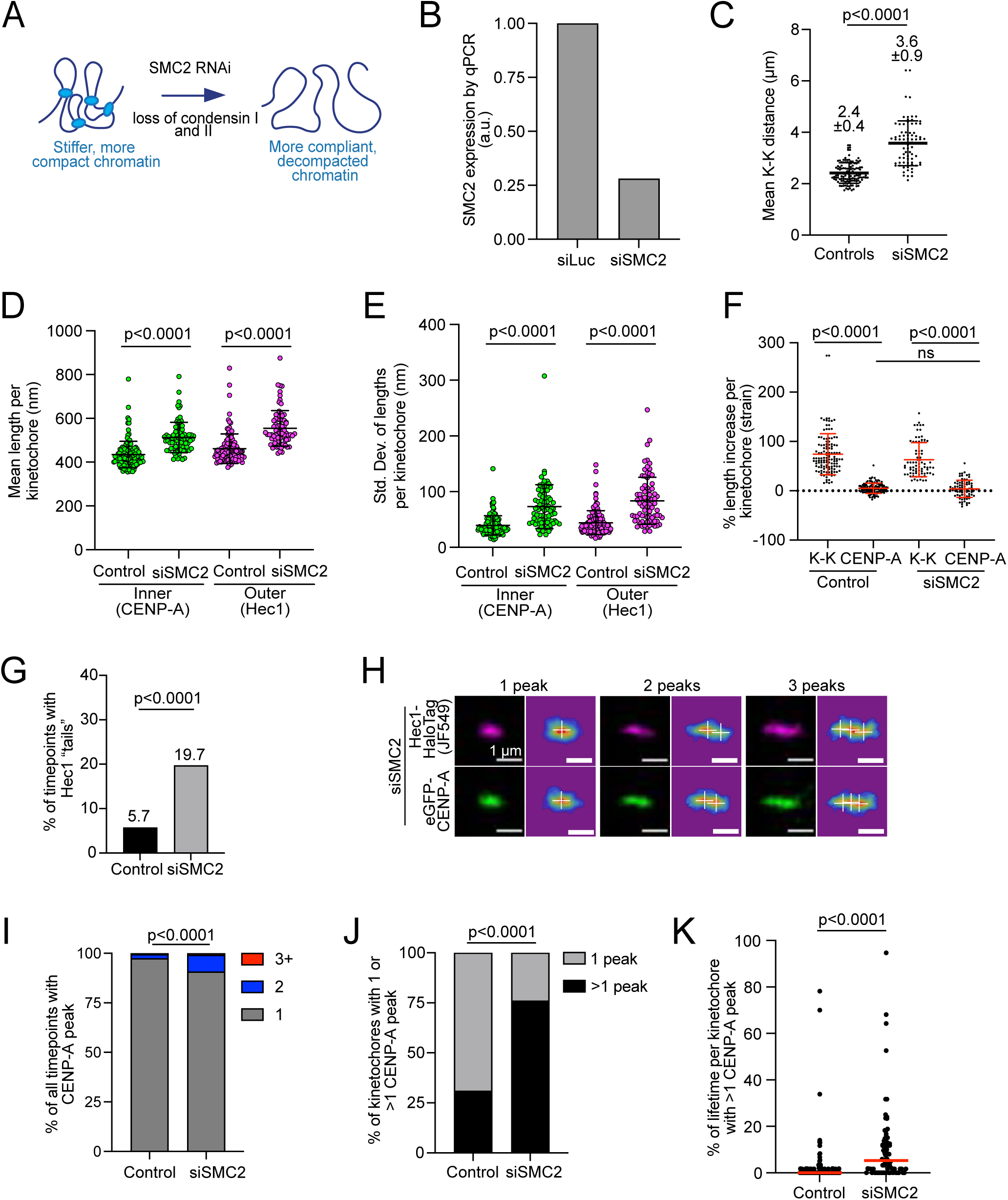
**Loss of condensin leads to variable metaphase kinetochore shape and loss of structural stability**. **(A)** Cartoon depicting loss of condensin I/II expecting to lead to more compliant chromatin. **(B)** Relative expression of SMC2 from control (siLuciferase) and siSMC2 cells from qPCR, normalized using actin and GAPDH as housekeeping genes. **(C)** Mean K-K distance per kinetochore pair for PtK2 control (m = 14 cells, n = 122 kinetochores) and SMC2 RNAi cells (m = 10 cells, n = 78 kinetochores) (mean ± SD; unpaired t-test). **(D)** Mean (from left to right: 435 ± 60 nm, 512 ± 70 nm, 461 ± 67 nm, 554 ± 81 nm) and **(E)** standard deviation (from left to right: 39 ± 17 nm, 73 ± 39 nm, 44 ± 22, 84 ± 42 nm) of length per kinetochore (eGFP-CENP-A and Hec1-Halotag with JF549 dye) over the imaging lifetime for control (n = 122 kinetochores) and siSMC2 cells (n = 78 kinetochores) (mean ± SD; Mann-Whitney test). **(F)** Percent of K-K distance and CENP-A length increase per kinetochore during imaging lifetime for control (n = 122; 74 ± 42% and 5 ± 10%) and siSMC2 cells (n = 78; 63 ± 35% and 4 ± 18%) (mean ± SD; Mann-Whitney test). **(G)** Percentage of timepoints of all imaged kinetochores displaying Hec1 “tails” for control (n = 6502 timepoints) and siSMC2 (n = 4075 timepoints) cells (Chi-square test). **(H)** Examples of Hec1 and CENP-A distributions with 1, 2, and 3 peaks with corresponding peak heatmap (multicolor images) and detection (white cross). **(I)** Percentage of timepoints of all imaged CENP-A kinetochores with 1, 2, or ≥3 peaks for control (n = 6502 total timepoints) and siSMC2 (n = 4075 total timepoints) cells (bottom) (Chi-square test). **(J)** Percentage of kinetochores for control (n = 122 kinetochores) and siSMC2 (n = 78 kinetochores) with >1 peak (31% for control and 76% for siSMC2) and with only 1 peak (69% for control and 24% for siSMC2) during its imaging lifetime (Chi-square test). **(K)** Percentage of each individual kinetochore’s imaging lifetime with >1 peak in Hec1 for control (n = 122 kinetochores) and siSMC2 (n = 78 kinetochores) cells (median; Mann-Whitney test).

We compared kinetochore lengths by live iSIM imaging, with and without SMC2 RNAi, in PtK2 cells (Fig. 4 D). For both eGFP-CENP-A and Hec1-HaloTag (+JF549), individual kinetochores in SMC2 RNAi cells were longer than control (Fig. 4 D), and had a higher standard deviation of lengths (Fig. 4 E). Thus, without SMC2 the kinetochore undergoes larger-scale deformations and actively changes shape more over time. We then asked how SMC2 RNAi impacts relative stiffness of the centromere and kinetochore, and calculated the approximate strain for each kinetochore and centromere using the percent increase of length as in control (Fig. 3 C). Similar to control kinetochores under endogenous spindle forces, SMC2 RNAi kinetochores also had a lower strain than SMC2 RNAi centromeres (Fig. 4 F), also consistent (Fig. 3 C) with kinetochore-specific elements stiffening the kinetochore.

Kinetochores in SMC2 RNAi cells were not only longer than in control cells but took on more complex and variable shapes. SMC2 RNAi kinetochores had more frequent Hec1 “tails” than control (Fig. 4 G), consistent with increased frequency of merotely with depleted condensin (Fig. 2) (Samoshkin et al., 2009). Further, SMC2 RNAi kinetochores displayed more frequent multimodal shapes along the force axis than control on a population-wide basis, with two or more intensity peaks detected in the summed 2D image intensity linescans (Figs. 4 H and I; and Video 4). More kinetochores overall exhibited intensity peaks (Fig. 4 J) and had more than one intensity peak during their imaging lifetime with SMC2 RNAi compared to control (Figs. 4 K and S3 A; and Video 4). This is consistent with centromeric and inner kinetochore organization in multiple partitions (Kixmoeller et al., 2025; Sacristan et al., 2024). While SMC2 RNAi kinetochores did not have peaks with significantly longer durations than control (Fig. S3 C), they visited multimodal states more frequently than control (Fig. 4 K; Figs. S3 A, B, and C; and Video 4). Together, we conclude that chromatin compaction, regulated by condensin I/II, is needed for structural homogeneity between kinetochores and structural maintenance over time within a kinetochore.

### Loss of condensin results in impaired kinetochore structure and function, but not elasticity, under acute forces

Given that kinetochores take on more extended and varied shapes with reduced SMC2 (Fig. 4), we used microneedle manipulation to probe the role of SMC2 in kinetochore material properties. We used PtK2 cells stably expressing eGFP-CENP-A, added SiR-tubulin dye and treated them with SMC2 siRNA for 48 h. To probe elastic vs plastic material properties in kinetochores with reduced SMC2, we looked at kinetochores that had a deformation of >100 nm with the microneedle pull. We measured CENP-A length before pulling, at maximum measured K-K, and after microneedle pulling, and pulled with the same parameters as control (Fig. 3). After pulling, most kinetochores relaxed to their baseline length (Figs. 5 A, B, C, and D; and Video 5), suggesting global maintenance of kinetochore elasticity despite condensin loss. As in control cells (Fig. 3 L), kinetochore and centromere strains were indistinguishable under exogenous force in SMC2 RNAi cells (Fig. 5 E). However, 2/9 siSMC2 kinetochores had dramatic tail lengthening at their maximum K-K distance that was persistent for greater than 30 s during the relaxation period (Figs. 5 F and G; and Video 6) compared to 0/6 control pulls that instead had broader, homogenous deformations. This suggests that condensin loss leads to impaired structural organization and maintenance under high force. We conclude that under high force, condensin is essential for kinetochores’ *local* structural organization but not for their *global* elastic properties.

**Figure 5:**
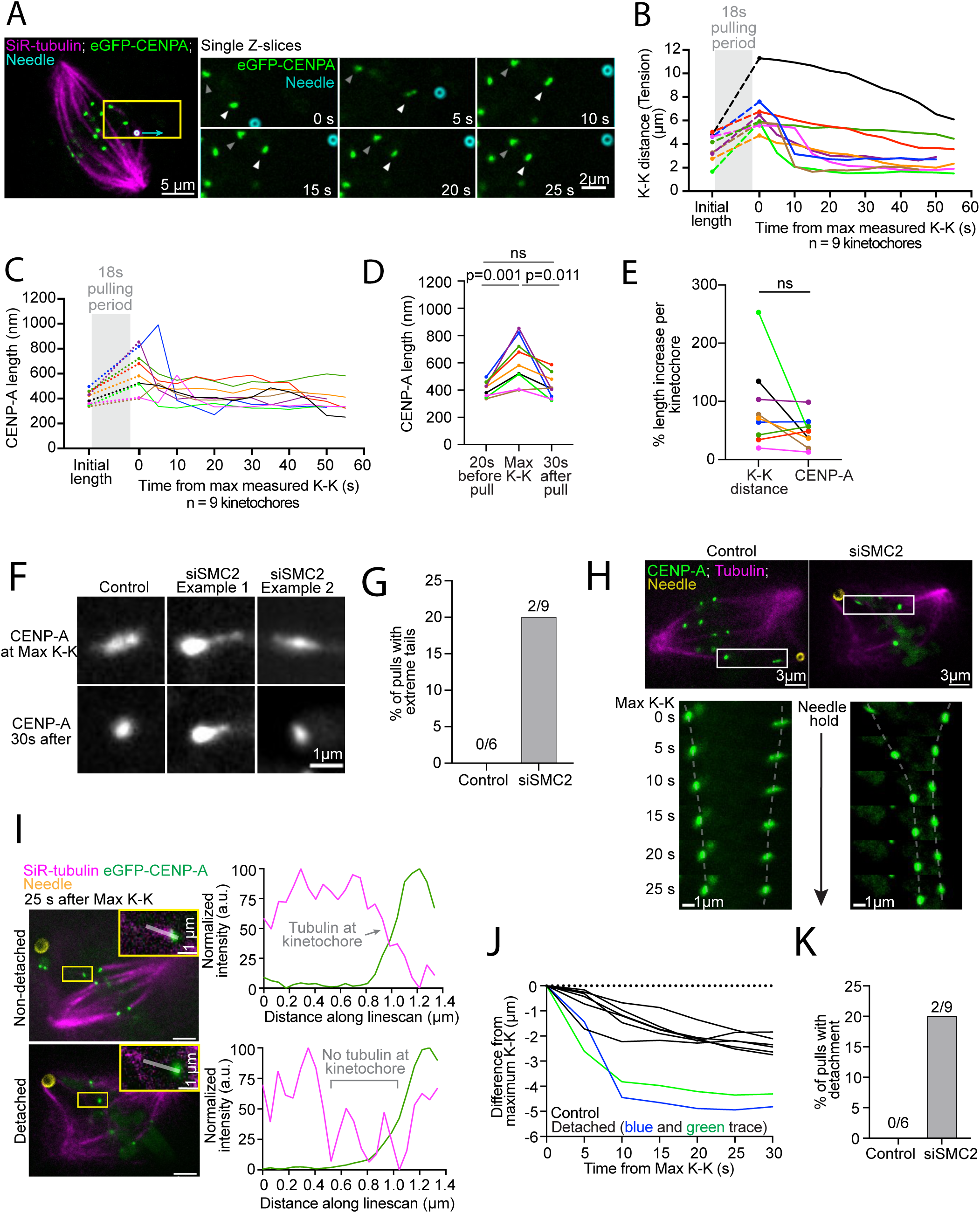
Loss of condensin results in impaired kinetochore structure and function, but not elasticity, under acute forces. **(A)** Representative PtK2 SMC2 RNAi cell timelapse of eGFP-CENP-A deforming with the microneedle pulling, and relaxing. Yellow box shows the zoomed in pair on the right, cyan arrow shows direction of needle movement, white arrowheads highlight the “front” kinetochore, and grey arrowheads highlight the “back” kinetochore. **(B)** K-K distance over time before, during, and after microneedle pulling for n = 9 kinetochores. “Initial” length is the frame just before pulling begins. Time = 0 s corresponds to the time maximum K-K distance was reached and measured during pulling, highlighting relaxation dynamics of each pulled kinetochore. The grey box indicates the pulling period. **(C)** CENP-A lengths over time before, during, and after microneedle pulling for 9 kinetochores as defined in (B). **(D)** CENP-A length at the frame before microneedle pulling, at the maximum K-K distance during pulling, and 30 s after pulling in siSMC2 cells (n = 9 kinetochores; Paired t-test). **(E)** Percent of K-K distance and CENP-A length increase from the frame before pulling begins to maximum measured K-K distance (n = 9 kinetochores; Paired t-test). **(F)** eGFP-CENP-A images at maximum measured K-K distance and 30s afterwards of control deformation and two siSMC2 kinetochore deformations exhibiting dramatic “tails”. **(G)** Percentage of control (n = 6) and siSMC2 (n = 9) kinetochore pulls with persistent “tails” for > 30 s during the relaxation period. **(H)** Timelapse comparing representative K-K distance relaxation for a control pull and for a fast relaxation SMC2 RNAi pull during the needle hold. White box indicates zoomed in pair. Grey dashed lines project the kinetochores’ movements over time. **(I)** Examples of non-detached (15 s after maximum K-K) and detached kinetochore (10 s after maximum K-K) from siSMC2 pulling experiments with fast K-K relaxation rates. Inset is of the front kinetochore with a linescan for tubulin intensity and kinetochore intensity; corresponding plots on the right annotated with sections of intensity signal corresponding to k-fiber presence or not. **(J)** K-K distance change after maximum K-K distance (t = 0) for control (black, n = 6 kinetochores), and detached, siSMC2 kinetochores (red and brown line, n = 2/9 kinetochores) pulls. **(K)** Percentage of pulls that led to detachment events in control and siSMC2 cells based on two criteria: fast K-K distance relaxation as in (J) and loss of tubulin signal attached to the kinetochore as in (I).

Unexpectedly, some SMC2 RNAi kinetochores appeared to detach from their k-fibers upon microneedle pulling. In total, 2/9 kinetochores had a dramatically faster K-K distance relaxation rate (from maximum K-K distance) compared control pulls (Figs. 5 H and J; and Video 3 and 7). Based on our previous work (Rosas-Salvans et al., 2025), we reasoned that kinetochore-microtubule detachment could cause such fast relaxation. Consistent with detachment, these same 2/9 kinetochores had no detectable k-fiber attached to them (Figs. 5 I and K; and Video 7). In contrast, we never observed detachment upon microneedle pulling in control cells herein (Fig. 3) or in previous work (Long et al., 2020; Rosas-Salvans et al., 2025; Suresh et al., 2020). Consistent with condensin depletion affecting CENP-A loading (Samoshkin et al., 2009; Yong-Gonzalez et al., 2007), we found a ∼20% decrease in Hec1 kinetochore intensity in SMC2 RNAi cells (Fig. S4 A). However, Aurora B kinase activity is known to decrease with condensin knockdown (Samoshkin et al. 2009), which would typically result in stronger kinetochore-microtubule attachments. Still, not just kinetochore structural instability but also kinetochore composition defects could contribute to increased detachment under reduced condensin.

Nevertheless, it is clear that kinetochore structural organization and attachment function are both compromised under high load with reduced condensin. Together, we conclude that the mammalian metaphase kinetochore requires condensin for robust material properties, structural organization, and function under high loads.

## Discussion

To perform their functions, kinetochores must both transmit and resist forces of different timescales, magnitudes and directions. Here, we ask how the mammalian kinetochore’s structure responds to and maintains itself under these forces. We find that inner and outer kinetochores together actively undergo large (1 µm) shape changes at metaphase (Fig. 1), and take on complex, irregular shapes under altered attachment geometries (Fig. 2). We demonstrate that metaphase kinetochores are elastic materials, deforming and recovering their shapes under endogenous and exogenous forces of different magnitudes and timescales (Fig. 3), and that their structural stability, homogeneity and resistance to microtubule detachment under force all depend on condensin (Fig. 4 and Fig. 5). Together, the work suggests that lateral sliding of individual protein linkages within the kinetochore mediates large scale shape changes and elastic material properties, with a centromere base that both allows and limits such sliding. We propose that the metaphase kinetochore has a flexible global structure to accommodate different force contexts without fracture or loss of function, and stiff local protein linkages to allow load-bearing and force transduction (Fig. 6).

**Figure 6:**
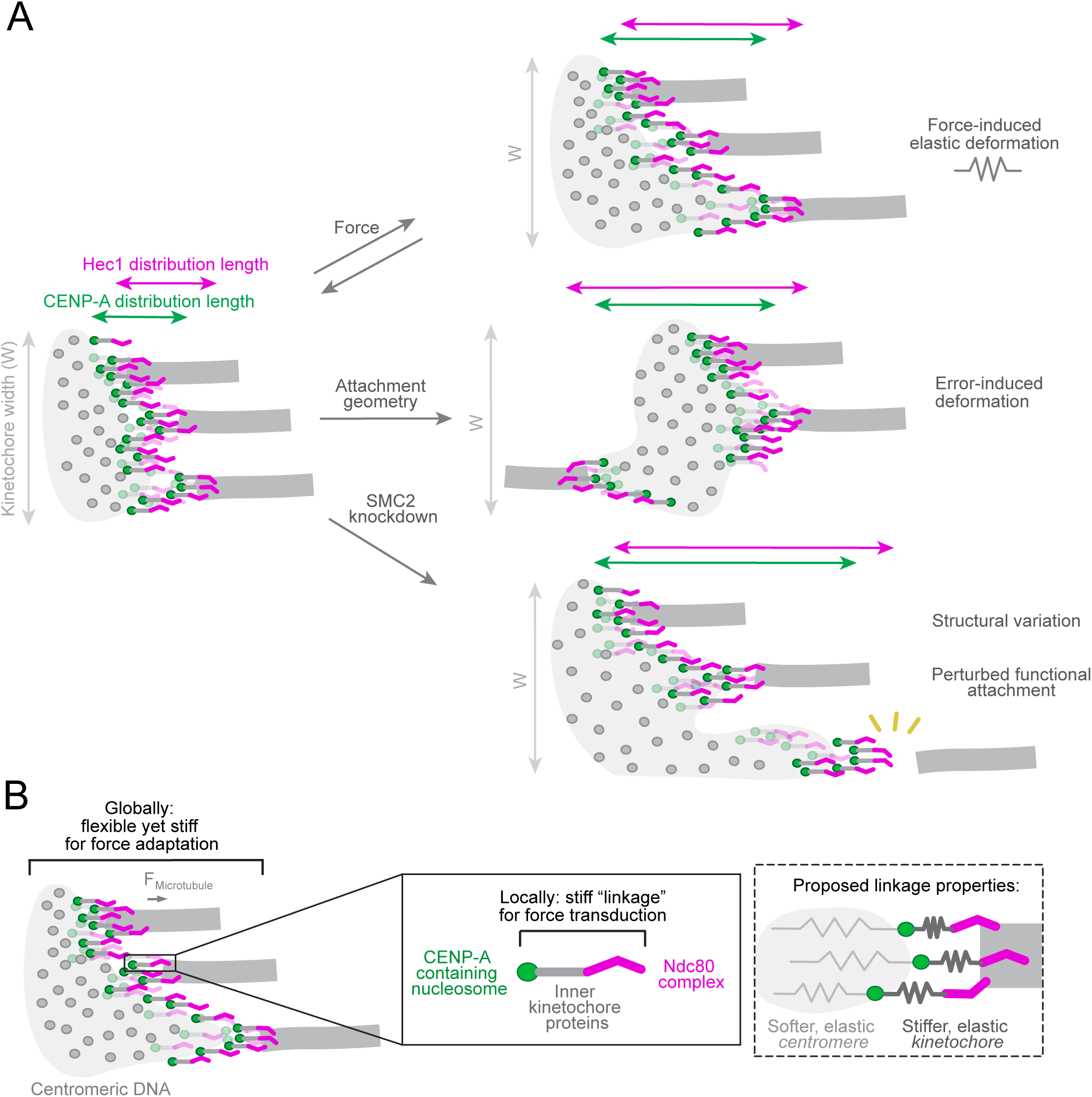
Model for mammalian metaphase kinetochore deformations and material properties. **(A)** Model of mammalian metaphase inner (green) and outer (magenta) kinetochore undergoing correlated, large-scale deformations in length (Fig. 1) by dynamic, relative sliding of parallel kinetochore protein linkages with respect to each other yet not changing kinetochore width (W). The metaphase kinetochore deforms elastically under force by reversibly sliding its protein linkages (top, Fig. 3), deforms under altered attachment geometry by sliding and reorienting linkages (middle, Fig. 2) and with reduced condensin slides linkages more variably and detaches from microtubules more (bottom, Figs. 4 and Fig. 5). **(B)** We propose that while individual kinetochore linkages are stiff enough (thick spring) to limit their deformation under load and allow local mechanotransduction, global metaphase kinetochore structure is compliant enough (thin spring) to adapt to varied attachment forces and geometries yet stiff enough to maintain attachment function. Zoom is of a local inner-to-outer kinetochore protein linkage that is stiff to allow for force transduction.

The model that parallel protein linkages slide with respect to each other to give rise to large-scale kinetochore shape deformations (Fig. 1 F and Fig. 6) raises the question of what drives such sliding. The force generated by individual microtubule plus-end tips within the k-fiber could in principle drive sliding. While the dynamics of plus-ends within a k-fiber must be somewhat coordinated to support chromosome movement over micrometers, not all tips are well aligned (Kiewisz et al., 2022; Leeds et al., 2023; O’Toole et al., 2020) or even in a single bundle (Kiewisz et al., 2022; Sacristan et al., 2024) when bound to the kinetochore. Accommodating offset tip positions would require that the protein linkages bound to microtubules are also offset. Supporting microtubule forces driving sliding, inner and outer kinetochore widths do not correlate with kinetochore length increases (Fig. S1 C and D), suggesting that large-scale length changes are due to inter-linkage sliding specifically in the microtubule force axis. Erroneous attachments, where microtubule tips take on not only offset positions but a different geometry, would require both the sliding of parallel protein linkages and pivoting of linkages to bind microtubules facing the opposite spindle pole. Consistently, erroneous attachments result in even more varied kinetochore shapes (Fig. 2 F and G) (Cimini et al., 2001; Khodjakov et al., 1997). How the material properties of the kinetochore, and inter-linkage sliding in our model, impact the coordination of microtubule dynamics within the k-fiber and error correction remains an open question.

While centromeres were proposed to be plastic (Lončarek et al., 2007) and merotelic anaphase kinetochores viscoelastic with a plastic component (Cojoc et al., 2016), we find that the metaphase kinetochore is an elastic material with no detectable plastic component, even under SMC2 RNAi (Fig. 3 C and I; Fig. 5 C and D). The origin of this elasticity is not known. In principle, it could come from elastic centromere material properties. Indeed, SMC2 RNAi impacts kinetochore structure (Fig. 4), and chromatin stiffness and organization have been found to influence the kinetochore’s structural organization (Ono et al., 2004; Sacristan et al., 2024; Samoshkin et al., 2009; Zielinska et al., 2019). Since condensin compacts chromatin (Hudson et al., 2003; Ono et al., 2003; Sun et al., 2018), condensin can impact how kinetochore proteins are assembled and interact with each other, e.g. establishing centromeric bipartite structure (Sacristan et al., 2024). However, and to our surprise, we found that condensin is not essential for kinetochore elasticity (Fig. 5 C and D). Whether chromatin-specific mechanisms, such as cohesins (Michaelis et al., 1997; Stephens et al., 2011), or kinetochore-specific mechanisms, such as oligomerization of kinetochore proteins (Hara et al., 2023; Sissoko et al., 2024) or presence of lateral crosslinkers (McKinley et al., 2015; Zhou et al., 2022), impact global kinetochore elasticity remains an open question. The fact that the centromere appears softer than the kinetochore (Fig. 3 C; Fig. 4 F) suggests that there must be some kinetochore-specific mechanisms. Notably, since kinetochores do not deform under high exogenous force along the chromosome arm axis (Biggs et al., 2025), but deform under endogenous and exogenous force along the microtubule axis (Fig. 1 E; Fig. 3 D), kinetochore material properties are likely direction dependent.

Centromere and kinetochore elasticity, and the difference between them, are likely important for function. If the centromere were too soft, this could impact force transmission to the sister kinetochore (Stephens et al., 2011), kinase signaling (Samoshkin et al., 2009), and coordination of microtubule tip dynamics (Leeds et al., 2023). A softer centromere under SMC2 RNAi (Fig. 4 F) could, through any of the above, lead to the loss of kinetochore grip robustness under force we observe (Fig. 5 K). Alternatively, centromere decompaction under SMC2 RNAi could lead to lower density of outer kinetochore microtubule binders or decreased binding between putative lateral crosslinkers reinforcing the kinetochore (McKinley et al., 2015; Rago & Cheeseman, 2013). In turn, if the centromere were too stiff, this could lead to a kinetochore that is too stiff and has decreased flexibility to adapt to different force contexts, attachment geometries, and microtubule tip positions. In addition, a too stiff centromere would require both sister kinetochore to have perfect plus-end dynamics coordination (of polymerization and depolymerization) for bioriented chromosome movement, and initial microtubule capture during spindle assembly may be less efficient due to limited conformational flexibility. Kinetochore elasticity could also directly contribute function. An elastic kinetochore could fully relax (Fig. 3 I and Fig. 5 D) after incorrect attachment to promote correct attachment formation or after being deformed by uncoordinated microtubule tips. Key to our model, the kinetochore being stiffer and centromere softer would allow the kinetochore to bind microtubules in diverse configurations while keeping force-transducing CENP-A-to-Hec1 linkages structurally intact – and thus functionally intact for mechanotransduction.

The mammalian metaphase kinetochore is a fatigue and fracture resistant material (Meyers et al., 2008; Qiang et al., 2019; Suresh, 1998; Vincent, 1982): it endures repeated stress cycles, and undergoes high strain (Fig. 1 E) yet does not fracture under endogenous cyclic loads (Fig. 1 C) or high exogenous loads (Fig. 3 I and Fig. 5 D). This parallels certain materials made by engineers (Li et al., 2022), with structural robustness coming from the material’s ability to absorb energy by deforming, to maintain load-bearing ability and to elastically recover after load application (Fig. 6). Of particular interest is the question of how kinetochore material properties – and underlying mechanisms – change to accommodate higher numbers of bound microtubules across species, including the dramatic example of holocentric kinetochores (Drinnenberg & Akiyoshi, 2017). With more microtubules bound to a kinetochore may come the need to both allow inter-linkage sliding to accommodate different tip positions and to laterally reinforce the kinetochore to limit such sliding. Determining the mechanisms that give rise to the kinetochore’s material properties and emergent mechanics, and how these change during mitosis and across evolution will be key to understanding the kinetochore’s physical functions.

## Acknowledgements

We thank Daniela Cimini for the rat kangaroo CENP-A construct, Dyche Mullins for providing the iSIM, Sam Lord for help optimizing iSIM parameters and with deconvolution. We thank Aussie Suzuki, Tanner Fadero, Wallace Marshall, Geeta Narlikar, and members of the S. Dumont laboratory for helpful discussions. This work was supported by the National Institutes of Health (NIH) R35GM136420, National Science Foundation Center for Cellular Construction DBI-1548297, NIH T32GM007810 (V.M. Tran), NIH T32GM139786 (V.M. Tran), NIH T32EB009383 (V.M. Tran), NIH F31GM156031 (V.M. Tran), NSF Graduate Research Fellowship Program (C.J.R.), UCSF Graduate Research Mentorship Fellowship (D.G.S.).

## Author contributions

V.M. Tran: Conceptualization, Data curation, Formal analysis, Funding acquisition, Investigation, Methodology, Project administration, Software, Supervision, Validation, Visualization, Writing—original draft, Writing—review and editing; J. Tao: Data curation, Formal analysis, Methodology, Software, Validation, Writing—review and editing; C.J. Rux: Investigation, Methodology, Writing—review and editing; D. Gomez Siu: Investigation, Methodology, Writing—review and editing; M. Rosas-Salvans: Conceptualization, Methodology; S. Dumont: Conceptualization, Funding acquisition, Project administration, Supervision, Writing—review and editing.

## Materials and Methods

### Cell culture, siRNA transfection, and protein expression

A PtK2 cell line stably expressing eGFP-CENP-A was generated from a wildtype PtK2 line (ATCC), infected with lentivirus for eGFP-CENP-A and selected with 0.5 µg/mL puromycin. Transient expression of Hec1-Halotag and Hec1_9A-Halotag constructs were done with lentiviral infection. Lentivirus was generated in HEK293T cells, and PtK2 cells were infected 48–72 h prior to imaging. All PtK2 cells were cultured at 37 °C and 5% CO2 in MEM (Invitrogen) supplemented with 5% sodium pyruvate (Invitrogen), 5% non-essential amino acids (Invitrogen), 5% penicillin/streptomycin (Invitrogen), and 10% qualified and heat-inactivated fetal bovine serum (Gibco). Cells were plated in 35 mm glass-bottom dishes (poly-D-lysine coated; MatTek Corporation) for live imaging experiments or in six-well plates with #1.5 25 mm coverslips (acid cleaned and poly-L-lysine coated) for immunofluorescence or without for immunoblotting. For RNAi experiments, siRNA targeting SMC2 (5′- AGTGTTGTAGTAGACACAGAG-3′) or luciferase (5’ CGUACGCGGAAUACUUCGA 3’, control) were transfected in the above Ptk2 eGFP-CENP-A cells using 4 µl of Oligofectamine (Life Technologies, 12252011) and 8 µL of 20 µM of siRNA in 288 µL Opti-MEM (Gibco), added to 1 mL of MEM + 10% FBS in imaging dishes. Five rat kangaroo SMC2 siRNA sequences were designed based on regions with highest homology to the human SMC2 sequence where existing SMC2 antibodies are successful. The above sequence was chosen based on its depletion efficiency and robustness judged qualitatively by microscopy in live-cells and verified by qPCR.

### Small molecule treatment

For live-cell experiments, microtubules were labeled by incubating cells with 100 nM SiR-tubulin and 10 µM verapamil (CY-SC002, Cytoskeleton Inc.) for a minimum of 30 min prior to imaging. Cells expressing HaloTag constructs were labeled with 200 nM JF549 (Promega) for a minimum of 30 min prior to imaging. For ZM447439 (Calbiochem) experiments, cells were regularly cycling and treated with 3 µM ZM447439 for a minimum of 30 min and a maximum of 90 min with the goal of imaging metaphase cells without severe kinetochore motility defects.

### Immunofluorescence

For immunofluorescence experiments quantifying Hec1 levels (Fig. S5 B), cells were plated onto acid-cleaned #1.5 25 mm coverslips coated with poly-L-lysine. Cells were fixed in 99.8% methanol for 3 min at -20 °C, 48 h after siRNA transfection, washed with TBST (0.05% Triton-X-100 in TBS), and blocked with 2% BSA in TBST (IF buffer hereafter). Antibodies were diluted in IF buffer and incubated for 1 h at room temperature (primary antibodies), washed 3x with TBST, then incubated for 45 min at room temperature (secondary antibodies). Cells were washed with TBST, then DNA was labeled with 1 mg/mL Hoescht 33342 for 1 min at room temperature prior to mounting on slides with Prolong Gold Antifade Mountant (P36934, Thermo Fisher). For immunofluorescence experiments quantifying tubulin levels closest to the kinetochore, cells were extracted in MTSB buffer (80 mM PIPES pH 6.8 + 1 mM MgCl2 + 1 mM EGTA + 0.5% Triton X-100) on ice for 1 min prior to methanol fixation as above, then processed as above. The following primary antibodies were used: mouse anti-Hec1 (1:1000; Novus Biologicals; NB100-338), chicken anti-GFP (1:500; Aves Lab Inc.; GFP-1010), rabbit anti-alpha tubulin (1:500; Abcam; ab18251), rat anti-alpha tubulin (1:2000; Bio-Rad; REFMCA77G). The following secondary antibodies were used at a 1:500 dilution in IF buffer: goat anti-chicken IgG Alexa Fluor 488 (Invitrogen; A-11039), goat anti-mouse IgG Alexa Fluor 568 (Invitrogen; A-11004), goat anti-rabbit IgG Alexa Fluor 405 (Invitrogen; A-31556), goat anti-rat IgG Alexa Fluor 647 (Invitrogen; A-21247).

### qPCR verification and analysis

PtK2 cells were seeded onto 6-well plates and transfected as above with a control Luciferase siRNA or a SMC2 siRNA for 48 h before cell collection for subsequent processing. Total RNA was extracted from the collected cells using the QIAwave RNA Mini Kit (Qiagen; 74534) following the manufacturer’s protocol. Cells were lysed using a 20 gauge needle and syringe and passed through 5-10 times. The Luna Universal One-Step RT-qPCR Kit (New England Biolabs; E3005L) was used with an input of 100 ng for each reaction. Both control and siSMC2 conditions were done with three biological replicates, each with technical triplicates. qPCR was run using kit instructions on the Biorad CFX Connect. Primer sequences for SMC2 (target), GAPDH (reference gene), and B-actin (reference gene) were generated and verified *in silico* based on the rat kangaroo transcriptome (Udy et al., 2015). They were then verified with gel electrophoresis using the Lunascript Kit (New England Biolabs; E3010L) to generate PtK2 cDNA from purified total RNA, and then performing PCR to check that the amplicon is of the expected size. The following primer sequences were used: SMC2 forward (F): 5ʹ-CCCAAGAGGAGCTGACCAAG -3ʹ and reverse (R): 5ʹ- GTGTTTCGCCACTTGTGCAT -3; B-actin (F): 5’-TGTACCCTCTCTCGGTCAGG-3’ and reverse (R): 5’-TGATGGTGTGACCCACACTG-3’; GAPDH (F): 5’-GTGTAGCCCAAGATGCCCTT-3’ and reverse (R): 5’- ACCTGAGCTGAATGGGAAGC-3’. qPCR analysis of knockdown was done using the Pfaffl method using multiple reference genes to normalize samples (Pfaffl, 2001).

### Microscopy and live-cell imaging

For all live cell experiments done without microneedle manipulation, live imaging was done using an inverted Nikon Ti-E equipped with a VT-iSIM super resolution module, 100-200 mW laser lines (405, 488, 561, and 642 nm), 100x 1.45 NA PlanApo oil objective (Nikon), Cairn Optospin emission filter wheel (ET 450/50m, ET 525/50m, ET 595/50m, ET 655lp, and ZET 405/488/651/640 m; Chroma Technology Corp.), and a Hamamatsu Quest interline-CCD camera (Fig. 1, 3, and 4). Cells were imaged with 11-13 z-planes 0.25 µm apart in 488 and 561 every 15 s, and a single central z-plane in 640 captured every 5 timepoints. After VT-iSIM imaging, images were deconvolved using Microvolution software (Microvolution). As such, object dimensions we measure are independent of imaging wavelength. As validation, we added TetraSpeck 200 nm Microspheres (Invitrogen) to the same imaging dishes with live cells and imaged regions with no cells but with beads in 488 nm and 561 nm wavelengths. Sizes were indistinguishable after deconvolution (Fig. 1 C and D). For live cell experiments with microneedle manipulation or for immunofluorescence imaging, imaging was done using an inverted Nikon Ti-E equipped with a spinning-disk confocal unit (CSU-X1; Yokogawa Electric Corporation) with Di01-T405/488/568/647 dichroic head (Semrock), 100x 1.45 Ph3 oil objective, 405 nm (100 mW), 488 nm (120 mW), 561 nm (150 mW), 642 nm (100 mW) diode lasers, emission filters (ET455/50M, ET525/50M, ET630/75M, and ET690/50M; Chroma Technology Corp.), and a Zyla 4.2 sCMOS camera (Andor Technology). For live experiments on the iSIM, images were collected at bin = 2 (92 nm/pixel) on Micro-Manager (2.0.0). For microneedle experiments and all immunofluorescence experiments, images were collected at bin = 1 (58 nm/pixel) on MicroManager (2.0.0). Cells were imaged in a humidified stage-top incubation chamber (Okolab) at 30 °C with 5% CO_2_ (Fig. 1, 2 and Fig. 4) or placed in CO_2_-independent media (Thermo Fisher) in a stage top chamber without the lid (Tokai Hit) with the sample temperature set at 30 °C (Fig. 3 and 5).

### Microneedle manipulation

Microneedle manipulation and needle fabrication was performed as outlined in previous work (Long et al., 2020; Suresh et al., 2020). Microneedles were made from glass capillaries with an inner and outer diameter of 1 mm and 0.58 mm respectively (1B100-4 or 1B100F-4, World Precision Instruments). Glass capillaries were pulled with micropipette puller (P-87, Sutter Instruments), bent and polished using a microforge (Narishige International) according to the same specifications, parameters, and geometries described earlier (Suresh et al., 2020). These parameters allowed for the needle to approach cells orthogonal to the imaging plane and to conduct manipulations without rupturing the cell (Suresh et al., 2020). Prior to imaging, microneedles were coated with BSA-Alexa-555 (Invitrogen, A34786) by soaking them in coating solution for 60 s. Coating solution was obtained by dissolving BSA-Alexa dye and Sodium Azide (Nacalai Tesque) in 0.1 M phosphate-buffered saline (PBS) at a final concentration of 0.02% and 3 mM, respectively (Sasaki et al., 2012). This coating solution allows needles to be visualized via fluorescence imaging, aiding in positioning of the needle along a single k-fiber, near to, but not touching, the kinetochore at a z-height appropriate for pulling on k-fibers without overly deforming the membrane.

Mitotic cells for microneedle manipulation were chosen based on the following criteria: spindles in metaphase, bipolar shape with both poles in the same focal plane, and expressing eGFP-CENP-A. In the case of condensin RNAi, cells with clear knockdown phenotypes were chosen. These phenotypes included: increased K-K distance, enlarged chromosomes, and wider metaphase plates (Ono et al., 2004; Ribeiro et al., 2009). To monitor inter-kinetochore distance and kinetochore deformation, we aimed to pull on single k-fibers close to the top of the cell with properly bioriented kinetochore pairs that could be tracked for the duration of the manipulation.

The micromanipulator was mounted to the microscope body and positioned above samples (Suresh et al., 2020). Manipulations were performed in 3D using a x-y-z stepper-motor micromanipulator (MP-225, Sutter Instruments). A 3-axis-knob (ROE-200, Sutter Instruments) was connected to the manipulator via a controller box (MPC-200, Sutter Instruments). Prior to manipulation, the needle was positioned via phase contrast imaging at the approximate x-y position of the intended manipulation site ∼ 10 µm above the cell in z. While imaging every 5 s at a single z-plane the needle was manually lowered into place along a single k-fiber, near to, but not touching the kinetochore. If necessary, the needle’s position in x-y was changed by raising the needle above the cell and repositioned in x-y before approaching the intended k-fiber and adjoining kinetochore pair. Once properly positioned, imaging parameters were changed to include 5 z-planes (± 0.6 μm, 0.3 μm spacing) in only the kinetochore channel. All other channels were imaged at only the middle z-plane. Prior to initiating pulling, control kinetochores were imaged for 2 min to establish the baseline kinetochore shape and oscillation pattern. Upon completion of the 2 min period with the needle adjacent to, but not pulling on the k-fiber, a Python script (Suresh et al., 2020) was used to communicate with the Sutter Multi-link software (Multi-Link, Sutter Instruments) to move the microneedle 10 µm over 18 s (62.5 nm steps) approximately orthogonal to the sister kinetochore-to-kinetochore axis. This computer-controlled movement of the microneedle allowed for consistent, reproducible experiments across all studied conditions. Upon completion of needle movement, cells were imaged and the needle remained in place until the stretched kinetochore pair began to relax and normal oscillations continued.

To be analyzed, the manipulated cells had to demonstrate all listed attributes: cell health was not significantly impacted by manipulation (no cell rupture or significant damage to the spindle), the mechanically challenged kinetochore pair was clearly visible throughout the entirety of manipulation and was attached in a manner consistent with biorientation, the inter-kinetochore distance of the mechanically challenged pair increased during manipulation (indicating force transmission to the kinetochores), the needle never directly contacted a perturbed kinetochore pair, the cell displayed bright enough kinetochore and tubulin signals to perform reliable kinetochore shape analyses and verifiably pull on the k-fiber of a chosen kinetochore pair and positively identify kinetochore-microtubule detachments. To rule out the possibility of k-fiber fracture as opposed to detachment from the kinetochore, tubulin intensity near the suspected detachment site at the kinetochore-microtubule interface was examined. Potential motion blur during kinetochore movement in the relaxation period would be persistent in all analyzed frames and is estimated to be less than the change in length detected, suggesting changes in length are not due to this.

### Image analysis

#### Kinetochore shape analysis

For live cell timelapse imaging, kinetochores were aligned along the kinetochore to sister kinetochore (K-K) axis (Fig. 1-5). Double fluorescence channel kinetochore images were split in Fiji (Schindelin et al., 2012), and then summed for both channels. Kinetochores were tracked using the TrackMate plugin in Fiji (Tinevez et al., 2017), employing the Laplacian of Gaussian (LoG) detector with an estimated spot diameter of 1 µm to fully capture potentially deformed kinetochores. The intensity threshold was manually adjusted to exclude background noise. The Advanced Kalman tracker in TrackMate generated kinetochore trajectories, with the maximum linking distance set to 1 µm to capture oscillatory movements. Sister kinetochores were manually paired by confirming their positions and periodic oscillations along the pole-to-pole axis.

For each identifiable kinetochore track, a 42 × 42 pixel region of interest (ROI) centered on the kinetochore’s coordinates was cropped from the image stack at each time point. Images were rotated to align the x-axis with the kinetochore-to-kinetochore (K-K) axis and flipped to position the k-fiber on the left and DNA on the right. For shape analysis, the three centermost z-slices, selected based on the highest intensity in the z-stack, were summed along the z-axis.

Kinetochore masks were generated with a Max Entropy threshold elevated by seven times the background intensity standard deviation to suppress noise. The largest connected component (> 9 pixels) nearest to the reported kinetochore position was retained, and its boundary was dilated by three pixels to fully capture the kinetochore perimeter.

To exclude overlapping kinetochores during mask detection, for each kinetochore, the total intensities within its mask across all time points were collected and divided into high- and low-intensity groups using Otsu’s method. If the mean intensity of the high-intensity group exceeded 1.7 times that of the low-intensity group and the mean pixel count of the high-intensity mask exceeded 60 pixels, the high-intensity group was discarded, as it likely resulted from two kinetochores overlapping. The dilated mask was projected onto the x- and y-axes to obtain one-dimensional line scans. Along each axis, the mean background intensity was subtracted to correct for residual background and dilation effects. Kinetochore lengths (x-axis) and widths (y-axis) were quantified using the area under curve normalized by the highest intensity (AUC/Peak) which is mathematically equivalent to a generalization of full-width half-maximum (FWHM) i.e. measuring the edge-to-edge distance of line scan at every intensity level from 0% to 100% of the maximum and then averaged (Fig. 1 A). A comparison is shown in Fig. 1 B between measuring by Gaussian fit FWHM and AUC/Peak, where the fit was generated by the built-in 1D Gaussian fit in Fiji.

For kinetochore poleward (P) and anti-poleward (AP) movement analysis, we quantified kinetochore displacement between consecutive frames and projected this displacement onto the kinetochore-to-kinetochore axis (K–K axis). A net displacement directed away from the sister kinetochore was classified as poleward, whereas displacement toward the sister kinetochore was classified as anti-poleward. In Fig. S2 A, we highlight kinetochores that exhibited two consecutive poleward intervals followed by three consecutive anti-poleward intervals, demonstrating a transition in stable directional movement.

For asymmetry analysis, the area under the one-dimensional intensity profile was integrated separately on the k-fiber (left) and DNA (right) sides of the primary peak (Figs. 2 E and F). Under unsaturated imaging conditions, a kinetochore was deemed asymmetric if there was an unbalanced intensity distribution. To account for pixel-level uncertainty at the primary peak point, the boundary was shifted from the leftmost to the rightmost edge of the central pixel, generating upper and lower bounds for the integrated intensity on each side. Asymmetry was scored only if using both leftmost and rightmost edge gave a consistent result. Before integration, the linescan was low-pass filtered with a finite-impulse-response (FIR) kernel designed with a Hamming window (cut-off = 2 cycles pixel⁻¹) to suppress high-frequency noise. A kinetochore was classified as having an intensity “tail” when the first derivative of the smoothed profile contained at least three significant local extrema whose amplitudes exceeded the background mean by at least half of the background standard deviation (Figs. 2 E, G, H, and I; and Fig. 4 H). For persistent “tail” formation (Figs. 5 F and G), kinetochores must have a “tail” for greater than 30 s during the kinetochore’s relaxation period measured over 1 min. For intensity peak analysis in two-dimensional maps (Figs. 4 I, J, K, and L), a local maximum was accepted as a true intensity peak only when its signal exceeded both 10% of the global maximum and the kinetochore boundary intensity by at least three standard deviations of the background intensity, thereby eliminating noise-induced peaks.

#### Strain analysis

To calculate strain (Fig. 3 C and Fig. 4 G), the three highest K-K distance values in a kinetochore’s track were averaged, then the three lowest K-K distance values were averaged. The percent increase was calculated as the lowest average subtracted from the highest average, divided by the lowest average, then multiplied by 100. For kinetochore strain, the corresponding time frames from the three highest and lowest K-K distances were used to compute the percent increase using CENP-A length.

#### Detachment analysis

To determine loss of tubulin intensity by linescans (Fig. 5 I), the frame where K-K relaxation plateaus (10 s after the maximum K-K timeframe was analyzed with manual linescans drawn over the eGFP-CENP-A channel to denote where the kinetochore signal is and where there would be an expected loss of tubulin signal. Then, the same linescan’s intensity values for the SiR-tubulin signal was plotted to show loss of tubulin signal at similar low intensities as where the kinetochore signal is noted.

#### Immunofluorescence analysis

To compare Hec1 intensities in control siLuciferase and siSMC2 cells, a 1.1 µm diameter circle was drawn over every isolated kinetochore without any overlapping particles in a max intensity projection of the spindle and analyzed in FIJI, then three regions outside the spindle area were measured using the same circle size as background. The Hec1 intensity was calculated as (kinetochore integrated density) – (mean background integrated density).

### Statistical analysis

For all statistical tests comparing two groups that are non-Gaussian distributed, a Mann-Whitney test was used. For statistical tests comparing two groups that are Gaussian distributed, a paired or unpaired student’s t-test was used. For all correlation tests, a simple linear regression was done. For comparisons of groups with a categorical outcome, a Chi-square test (more than two variables) was used.

### Video Preparation

Videos were prepared using FIJI. Brightness and contrast were adjusted to clearly visualize the kinetochores and microtubules.

## Supplement Figure Legends

**Supplemental Figure 1:**
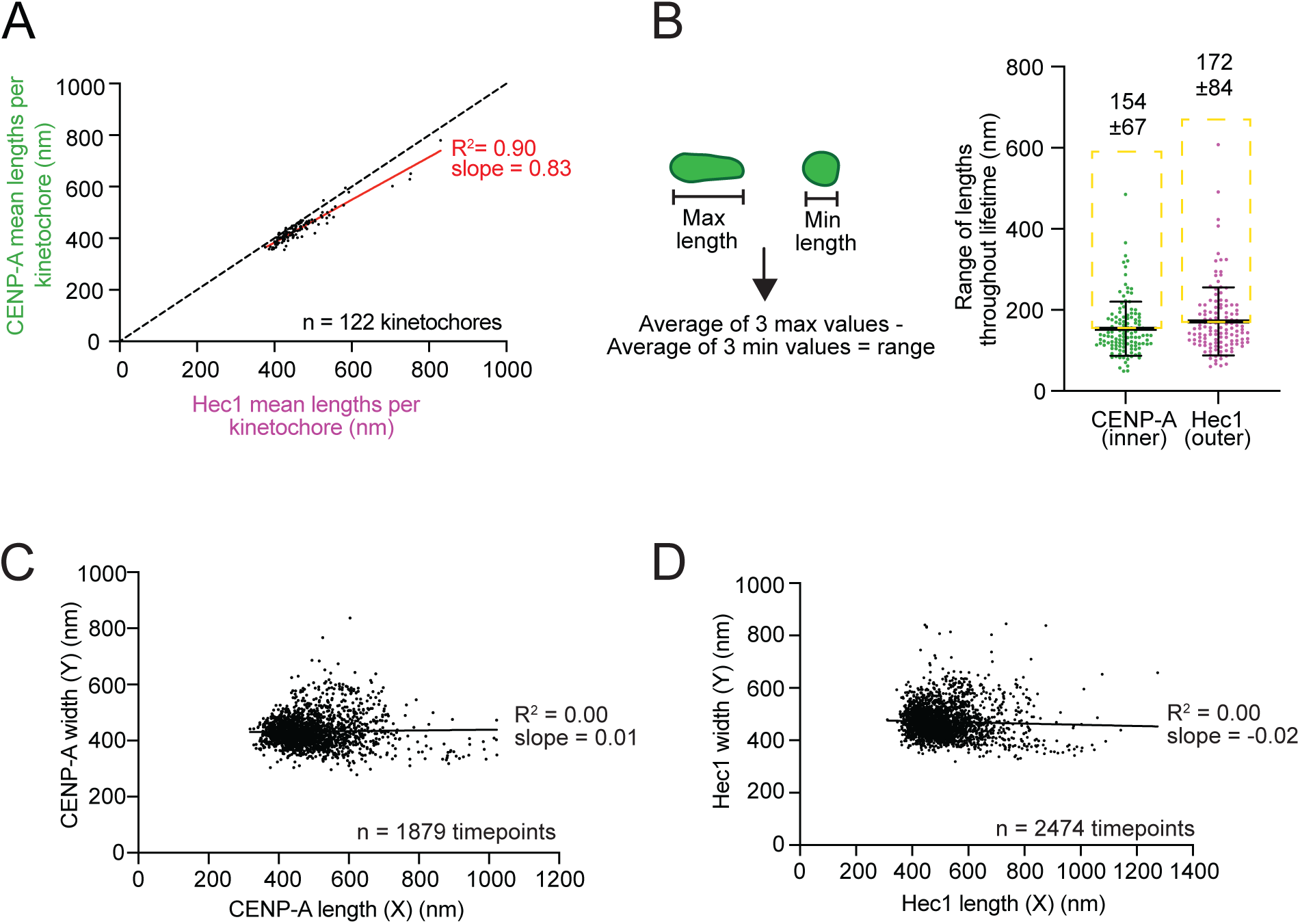
Inner and outer kinetochore lengths are correlated across individual kinetochores, and their widths do not correlate with length. (A) Correlation between CENP-A and Hec1 mean lengths for each kinetochore. Red line is the best linear fit and dashed black line is a 1:1 comparison (m = 14 cells, n = 122 kinetochores; R^2^ = 0.90; simple linear regression). (B) (Left) Cartoon of how the range of a kinetochore’s length throughout its imaging lifetime is calculated: the average of the three highest length values minus the average of the three lowest length values. (Right) Length range for all kinetochores, visualized by eGFP-CENP-A and Hec1-Halotag with JF549 dye with mean 154 ± 67 nm and 172 ± 84 nm, respectively (m = 14 cells; n = 122 kinetochores) and yellow dashed boxes denoting large deforming kinetochores (mean ± SD; Mann-Whitney test). (C) Correlation between CENP-A widths and lengths for large-deforming kinetochore timepoints as indicated in (B) with black line as the best linear fit (n = 1879 timepoints; R^2^ = 0.00; not significant from zero; Simple linear regression). (D) Correlation between Hec1 widths and lengths for large-deforming kinetochore timepoints as indicated in (B) with black line as the best linear fit (n = 2474 timepoints; R^2^ = 0.00; Significant from zero; Simple linear regression).

**Supplemental Figure 2:**
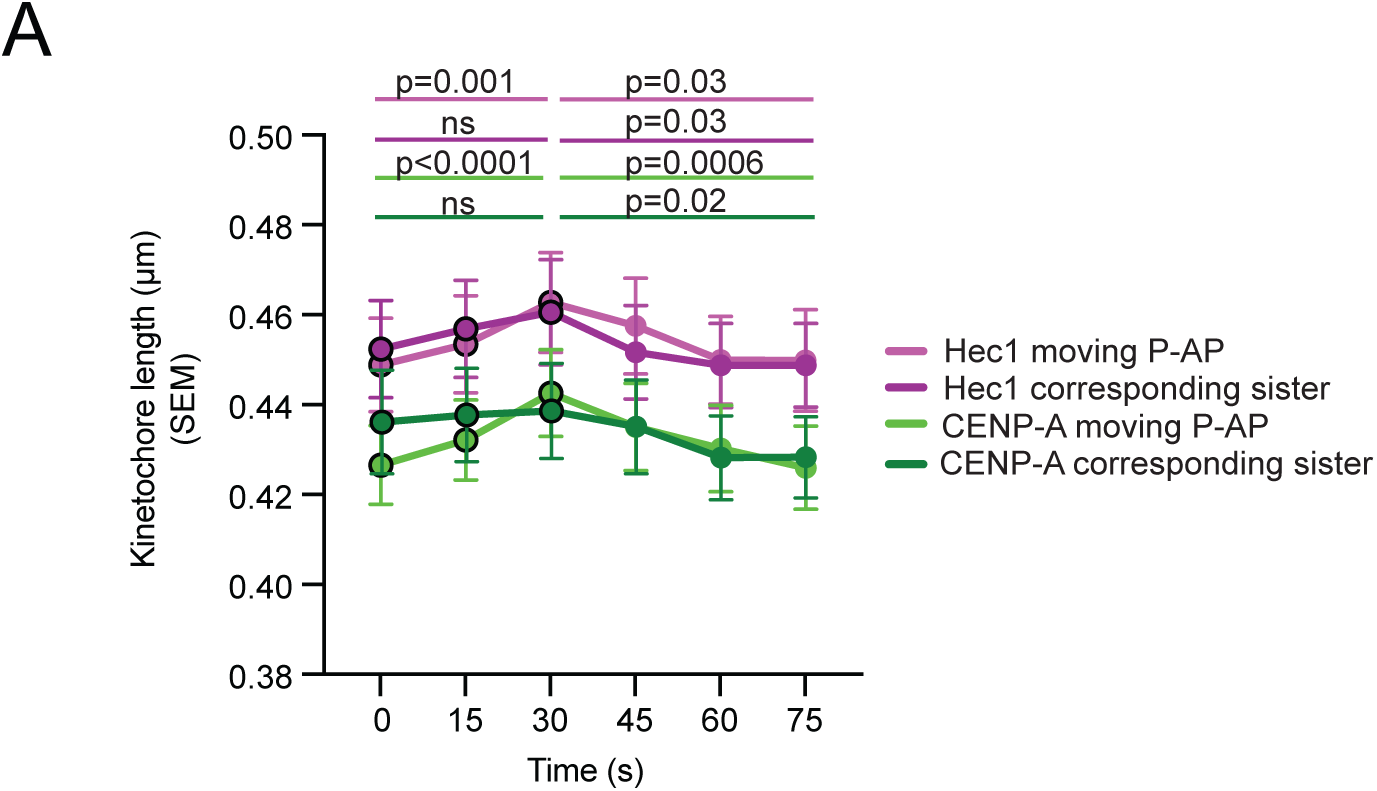
Inner and outer kinetochores both deform around kinetochore directional switches. (A) Plot of CENP-A and Hec1 lengths three timepoints before (0-30 s) and after (45-75 s) a directional switch (between 30-45 s) from poleward to antipoleward direction (P-AP) and their corresponding sister kinetochore lengths (m = 151 turnarounds; mean ± SEM; Mann-Whitney test).

**Supplemental Figure 3:**
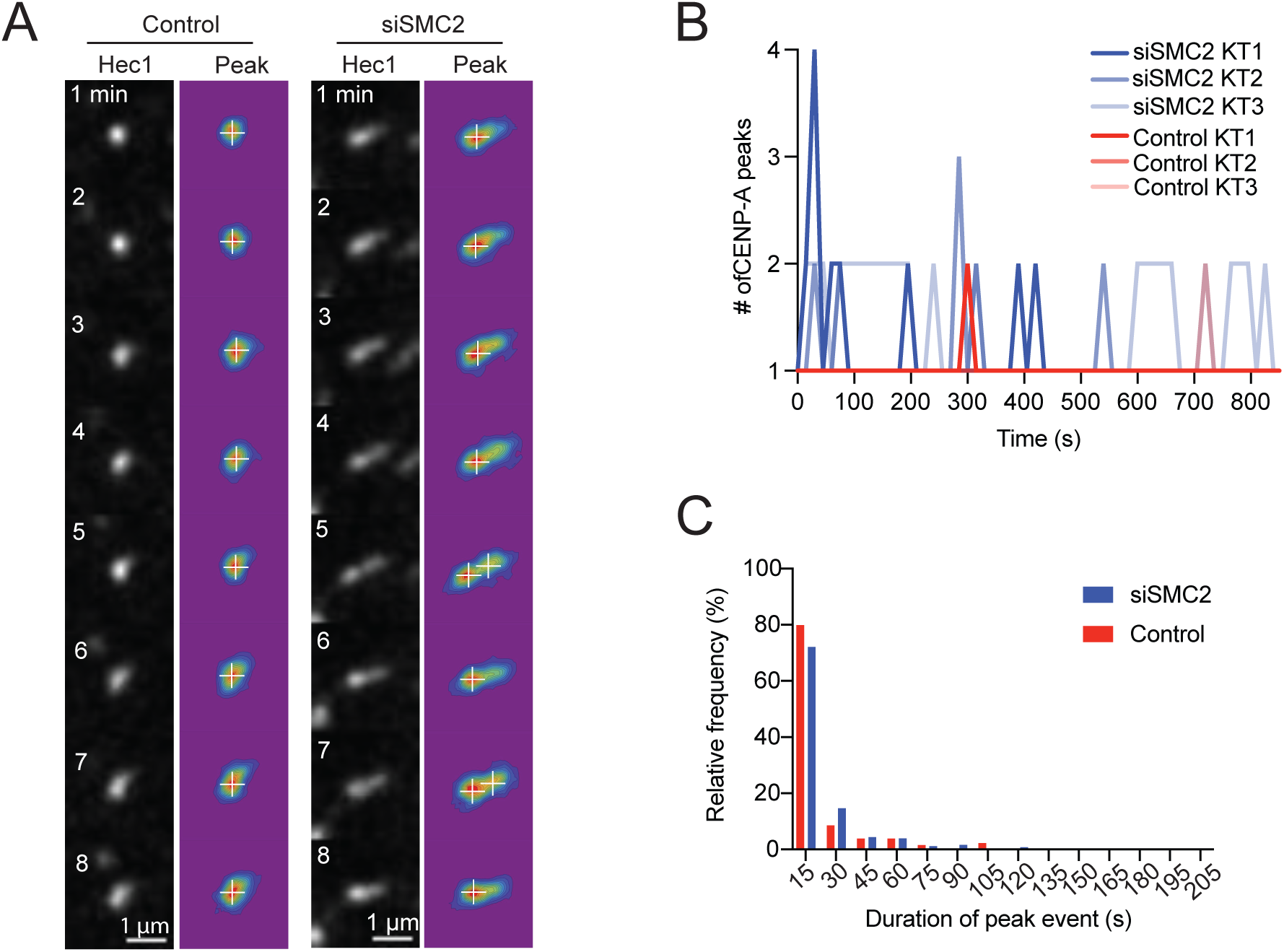
Loss of condensin results in more frequent multipeak kinetochore shapes but are not more long-lived. (A) Timelapse of control and siSMC2 kinetochore over time, visualizing Hec1-Halotag with JF549 dye and corresponding peak heat map (multicolor) and detection (white cross). (B) Example traces of three control (shades of red) and three siSMC2 (shades of blue) kinetochores and their CENP-A peaks detected over time. (C) Histogram of percentage of kinetochores with specified peak durations by frame (15 s) for control (n = 122) and siSMC2 (n = 78) kinetochores.

**Supplemental Figure 4:**
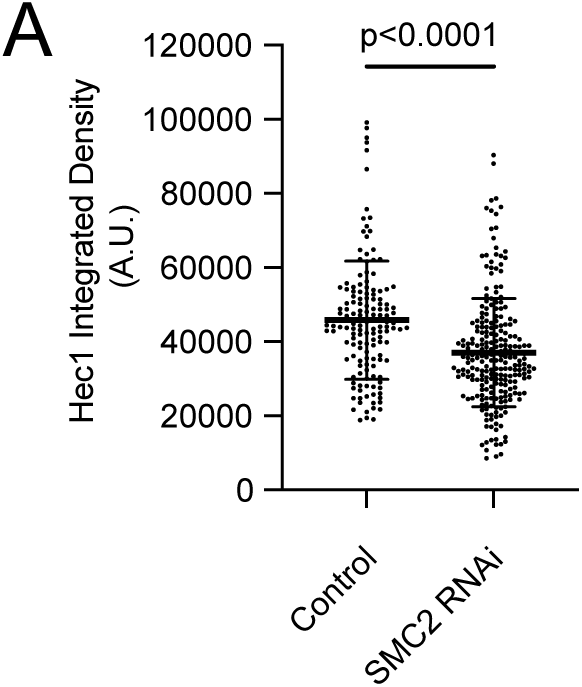
Kinetochores in SMC2 RNAi cells detach from microtubules under acute force application. (A) Hec1 kinetochore intensity from immunofluorescence comparing control siLuciferase (m = 10 cells; n = 147 kinetochores) and siSMC2 (m = 15 cells; n = 241 kinetochores) cells (mean ± SD; Unpaired t-test).

## Video legends

Video 1: Inner and outer kinetochores change shape at small and large scales during metaphase, related to Figure 1. Timelapse iSIM imaging (maximum-intensity projection) of a representative PtK2 control cell at metaphase visualizing eGFP-CENP-A and Hec1-Halotag with JF549 dye. Small-scale deforming kinetochore (white arrowheads) and large-scale deforming kinetochore (yellow arrowhead) is seen in metaphase. Shape analysis is done on the K-K axis (sister kinetochores indicated with open arrowhead). Eleven planes spaced 0.25 µm apart were acquired every 15 s. Scale bar, 5 µm. Time is in min:s. Playback speed is 10 frames/s. Video corresponds to still images from Fig. 1 B.

Video 2: Inner and outer metaphase kinetochores have larger, more complex deformations with Hec1-9A overexpression and ZM treatment compared to control, related to Figure 2. Timelapse iSIM imaging (maximum-intensity projection shown) of three PtK2 cells corresponding to control, overexpressed mutant Hec1-9A-Halotag, and 3 µM ZM447439 treatment for 30-90 min at metaphase. All conditions show eGFP-CENP-A and Hec1-Halotag with JF549 dye (or with Hec1-9A-Halotag). Small-scale deforming kinetochore pair for control cell (top cell, white arrowheads) is seen oscillating. Large-scale deforming kinetochore pairs (middle and bottom cell, white arrowheads) for 9A and ZM cells have dampened oscillations and show complex, asymmetric shapes. Zoomed in kinetochore pairs show eGFP-CENP-A signal and Hec1-Halotag with JF549 dye (or Hec1-9A-Halotag) separately. Eleven planes spaced 0.25 µm apart were acquired every 15 s. Scale bar in whole cell movies, 3 µm; in zoomed kinetochores, 1 µm. Time is in min:s. Playback speed is 10 frames/s. Video corresponds to kinetochore pairs shown in still images from Fig. 2 B.

Video 3: Metaphase CENP-A distribution deforms with microneedle pulling and relaxes back to baseline length, related to Figures 3 **and 5**. Timelapse spinning disk confocal imaging (single z-plane shown) of a PtK2 cell expressing eGFP-CENP-A and labeled with SiR-tubulin undergoing a microneedle pull of 10 µm over 18 s. Needle (pseudocolored yellow) is dyed with BSA-Alexa-555. The “front”, needle-proximal kinetochore (filled white arrowhead) deforms when pulled and is tracked with the “back” sister kinetochore (open white arrowhead). The needle pull and hold period is indicated in the top right. Five planes spaced 0.3 µm apart were acquired every 5 s during the pull and relaxation period. Scale bar is 3 µm. Time is in min:s. Playback speed is 2 frames/s. Video corresponds to still images in Fig. 3 F and Fig. 5 I.

Video 4: Inner and outer metaphase kinetochores under SMC2 RNAi have large deformations, exhibiting asymmetry and secondary peak formation, related to Figure 4. By end Timelapse iSIM imaging (maximum-intensity projection shown) of a PtK2 cell under SMC2 RNAi over 48 h, visualizing eGFP-CENP-A and Hec1-Halotag (+JF549 dye) in side-by-side movies. A kinetochore (white arrowhead) at metaphase has large-scale deformations (nearly 1 µm) and exhibits peak formation. Eleven planes spaced 0.25 µm apart were acquired every 15 s. Scale bar, 3 µm. Time is in min:s. Playback speed is 10 frames/s. Video corresponds to kinetochore shown in still images from Fig. 4 I and Figure S4 A.

Video 5: Metaphase CENP-A distribution in SMC2 RNAi cell deforms under microneedle pulling force, stretching and subsequently relaxing upon force removal, related to Figure 5. Timelapse spinning disk confocal imaging (single z-planes shown) of a PtK2 cell expressing eGFP-CENP-A and labeled with SiR-tubulin undergoing a microneedle pull of 10 µm over 18 s. Cell was under SMC2 RNAi over 48 h before imaging. Needle (pseudocolored yellow) is dyed with BSA-Alexa-555. The “front” needle-proximal kinetochore (white arrowhead) deforms during the pull and relaxes back to its baseline length. The needle pull and hold period is indicated in the top right. Five planes spaced 0.3 µm apart were acquired every 5 s before the pull, during the pull, and over the relaxation period. Scale bar is 3 µm. Time is in min:s. Playback speed is 2 frames/s. Video corresponds to still images in Fig. 5 A.

Video 6: Metaphase CENP-A distribution in SMC2 RNAi cell deforms with microneedle pulling force, revealing dramatic “tail” before relaxing upon force removal, related to Figure 5. Timelapse spinning disk confocal imaging (single z-plane shown) of a PtK2 cell expressing eGFP-CENP-A and labeled with SiR-tubulin undergoing a microneedle pull of 10 µm over 18 s. Cell was under SMC2 RNAi over 48 h before imaging. Needle (pseudocolored yellow) is dyed with BSA-Alexa-555. The “front” needle-proximal kinetochore (white arrowhead) deforms during the pull with a long and dramatic “tail”, reaching nearly 1 µm towards the chromatin. The sister kinetochore is visible in another plane. The needle pull and hold period is indicated in the top right. Five planes spaced 0.3 µm apart were acquired every 5 s during the pull and relaxation period. Scale bar is 3 µm. Time is in min:s. Playback speed is 2 frames/s. Video corresponds to still images in Fig. 5 G.

Video 7: Metaphase kinetochore in SMC2 RNAi cell deforms then detaches from k-fiber under microneedle pulling force, related to Figure 5. Timelapse spinning disk confocal imaging (single z-plane shown) of a PtK2 cell expressing eGFP-CENP-A and labeled with SiR-tubulin undergoing a microneedle pull of 10 µm over 18 s. Cell was under SMC2 RNAi for 48 h before imaging. Needle (pseudocolored yellow) is dyed with BSA-Alexa-555. The “front” needle-proximal kinetochore (white arrowhead) deforms, then detaches from its k-fiber as it moves back towards its sister. Bottom video is zoomed into the kinetochore with tubulin signal enhanced to better visualize detachment. White arrowhead follows the detaching kinetochore and a yellow arrowhead appears at the frame just before detachment from the k-fiber occurs. Five planes spaced 0.3 µm apart were acquired every 5 s during the pull and relaxation period. Scale bar is 3 µm. Time is in min:s. Playback speed is 2 frames/s. Video corresponds to still images in Fig. 5I and Fig. 5A.

